# A mucin-regulated adhesin determines the intestinal biogeography and inflammatory character of a bacterial symbiont

**DOI:** 10.1101/2022.07.25.501442

**Authors:** T. Jarrod Smith, Deepika Sundarraman, Ellie Melancon, Laura Desban, Raghuveer Parthasarathy, Karen Guillemin

**Author notes:** correspondence (KG).

## Abstract

In a healthy gut, microbes are often aggregated with host mucus, yet the molecular basis for this organization and its impact on intestinal health are unclear. Mucus is a viscous physical barrier separating resident microbes from epithelia, but also provides glycan cues that regulate microbial behaviors. Using experimental evolution, we discovered a mucin-sensing pathway in an *Aeromonas* symbiont of zebrafish, Aer01. In response to the mucin-associated glycan N-acetylglucosamine, a sensor kinase regulates expression of a mucin-binding adhesin we named MbpA. When MbpA is disrupted, Aer01 colonizes to normal levels, but is largely planktonic and elicits increased intestinal inflammation, traits which are reversed by increasing cell surface MpbA. MbpA-like adhesins are common in human-associated bacteria and expression of an *Akkermansia muciniphila* MbpA-like adhesin in MbpA-deficient Aer01 restored lumenal aggregation and reversed its pro-inflammatory character. Our work demonstrates how resident bacteria use mucin glycans to modulate behaviors congruent with host health.

## INTRODUCTION

Vertebrate intestinal mucus is vital for maintaining host-microbe mutualism. Mucus can modulate symbiosis by encapsulating and segregating resident microbes away from the gut epithelium (Johansson et al., 2008; Vaishnava et al., 2011). In a healthy digestive tract, microbes are often found associated with host mucus in the lumen, a distribution that has been described across vertebrates including humans, mice, and zebrafish (Bergstrom et al., 2020; Frese et al., 2013; Schlomann et al., 2018; Sonnenburg et al., 2005; Welch et al., 2017; Wiles et al., 2018). This lumenal microbial aggregation is commonly attributed to host processes such as active mucus secretion and peristalsis that encapsulate and displace lumenal contents, with the microbiota members being passive players within system. In this view, loss of gut health is attributed to host defects such as impaired mucus secretion or gut motility. Microbes however, can actively regulate their own aggregation and biofilm formation in response to environmental cues and recent studies indicate glycans associated with mucin can act as potent mediators of bacterial behaviors (Co et al., 2018; Hecht et al., 2017; Kavanaugh et al., 2014; Ottman et al., 2016; Pacheco et al., 2012; Wang et al., 2020; Wheeler et al., 2019). The dual roles of mucin as a physical barrier and molecular signal are inseparable since the sugars that decorate mucins contribute to both its biophysical properties and its chemical signature. Defining the ways in which microbiota members sense and respond to mucins to regulate their own structural organization in the gut will refine our understanding of microbiota roles on gut ecosystem health and disease.

Mucin is secreted by intestinal goblet cells, with *MUC2* being the dominant mucin in the human, mouse, and zebrafish gut (Jevtov et al., 2014; Massaquoi et al., 2022; Willms et al., 2021). Mucin is heavily glycosylated with O-linked N-acetylgalactosamine (GalNAc), galactose (Gal), and N-acetylglucosamine (GlcNAc) glycan chains that are usually capped with fucose (Fuc) and/or sialic acid (SA) (Bergstrom & Xia, 2022). The glycan composition of mucin varies along the intestinal tract (Larsson et al., 2009) and various microbes encode adhesins that bind glycan epitopes O-linked to mucin (Juge, 2012; Maresca et al., 2021). Resident microbes alter mucin’s glycan composition and stimulate its secretion (Arike et al., 2017; Bates et al., 2006; Bergstrom et al., 2020; Pickard et al., 2014). Additionally, microbes encode glycoside hydrolase (GH) enzymes that process glycan O-linkages on mucin, releasing energy-rich monomeric and multimeric glycan products that can also serve as molecular signals to microbiota (Crouch et al., 2020; Koropatkin et al., 2012; Ndeh & Gilbert, 2018; Wang et al., 2020; Wardman et al., 2022). The mechanisms through which mucin glycans may signal to microbiota members and whether these processes impact symbiosis remain to be explored.

Most studies exploring the relationship between mucin’s barrier function and microbial symbiosis use mouse models with complete loss or severe defects in intestinal mucin secretion and thus dramatic changes to the gut’s glycan landscape. Loss of mucin often results in increased microbial proximity to the epithelium associated with intestinal inflammation that can progress to various intestinal diseases such as colitis and colorectal cancer (Cullender et al., 2013; Johansson et al., 2011). Another feature of the mucus-depleted gut is elevated levels of bacterial flagellin, indicative of motile bacteria and also a driver of intestinal inflammation (Cullender et al., 2013; Hayashi et al., 2001). In these studies, intestinal mucus clearly promotes host health but the extent to which it does so as a barrier or through glycan-mediated regulation of microbial activities is difficult to disentangle.

The larval zebrafish has emerged as a powerful model system for dissecting mechanistic principles of vertebrate host-symbiont interactions due to its optical transparency, transgenic capabilities, relative ease of gnotobiology, and conservation of many mammalian innate immune processes (Burns & Guillemin, 2017; Flores et al., 2020). Live imaging of the larva zebrafish (*Danio rerio*) intestine demonstrates bacteria in the gut lumen adopt diverse aggregation states and localize to distinct regions of the gut (Schlomann et al., 2018; Wiles et al., 2018), similar to observations in fixed mouse intestinal tissue (Earle et al., 2015; Frese et al., 2013; Welch et al., 2017). For example, the zebrafish symbiont Aeromonas ZOR0001 (Aer01) forms dense aggregates in the anterior midgut with few planktonic cells, while the closely related *Aeromonas* ZOR0002 is more planktonic and located more anteriorly. Recent live imaging analysis and modeling of bacterial population dynamics has highlighted the importance of host processes, such as peristalsis (Wiles et al., 2016) and bacterial-regulated behaviors of motility (Wiles et al., 2020) and aggregate assembly and fragmentation as critical determinant of bacterial biogeography in the zebrafish gut (Schlomann et al., 2019; Schlomann & Parthasarathy, 2021).

Here, we use the Aer01-zebrafish mutualistic relationship as a model to investigate the relative contributions of mucin’s barrier and signaling roles in shaping symbiosis in a healthy host. We found Aer01 actively promotes host intestinal health by self-limiting its intestinal distribution and inflammatory character in response to the mucin-associated glycan GlcNAc. Experimental evolution that selected for “<u>m</u>ucin <u>b</u>lind” (MB) Aer01 isolates that do not aggregate in response to mucin or GlcNAc revealed that a two-component system (TCS) sensor histidine kinase (MbpS) and regulatory protein (MbpD) coordinate to control the transcription and function of an adhesin we called MbpA. Inhibition of MbpA function is sufficient to enhance intestinal inflammation and alter Aer01 population structure in the gut, indicating regulation of this adhesin in response to GlcNAc signals is crucial for modulating the inflammatory tone of the intestine. MbpA analogs are distributed in various human-associated microbiota including strains of *Oxalobacter, Fusobacterium, and Akkermansia*. Similar to MbpA, the MbpA analog in *Akkermansia* (Amuc_1620, here called MucA) is upregulated in the presence of mucin (Ottman et al., 2016). Expression of MucA in an Aer01 strain with a non-functional MbpA (MB1) rescued aggregation and suppressed its pro-inflammatory activity, suggesting MucA may play a similar role in beneficial *Akkermansia-human* interactions. Together, these data indicate Aer01 actively promotes its hosts’ intestinal health in response to the glycan landscape of the gut environment and suggests this principal may shape beneficial interactions in other host-microbe systems.

## RESULTS

### Aer01 is associated with host mucus in the intestinal lumen

To determine the spatial relationship between Aer01 and intestinal mucus, we mono-associated germfree larval zebrafish with dTomato (dTom)-tagged Aer01 four days post fertilization (dpf) and fixed whole larvae 24 hours later for confocal imaging. Aer01 distribution was visualized using anti-dTom and FITC-conjugated wheat germ agglutinin (WGA-FITC), which binds mucin-associated glycans including GlcNAc, was used as a proxy for mucus distribution. Consistent with previous histological studies of goblet cell distribution in the zebrafish intestine, we found two anatomically separated populations of mucus-producing goblet cells in the pharynx/esophagus and mid/distal gut indicated by WGA-positive vesicles (**Fig. 1A**) (Ng et al., 2005; Troll et al., 2018; Wallace et al., 2005).

**Figure 1.**
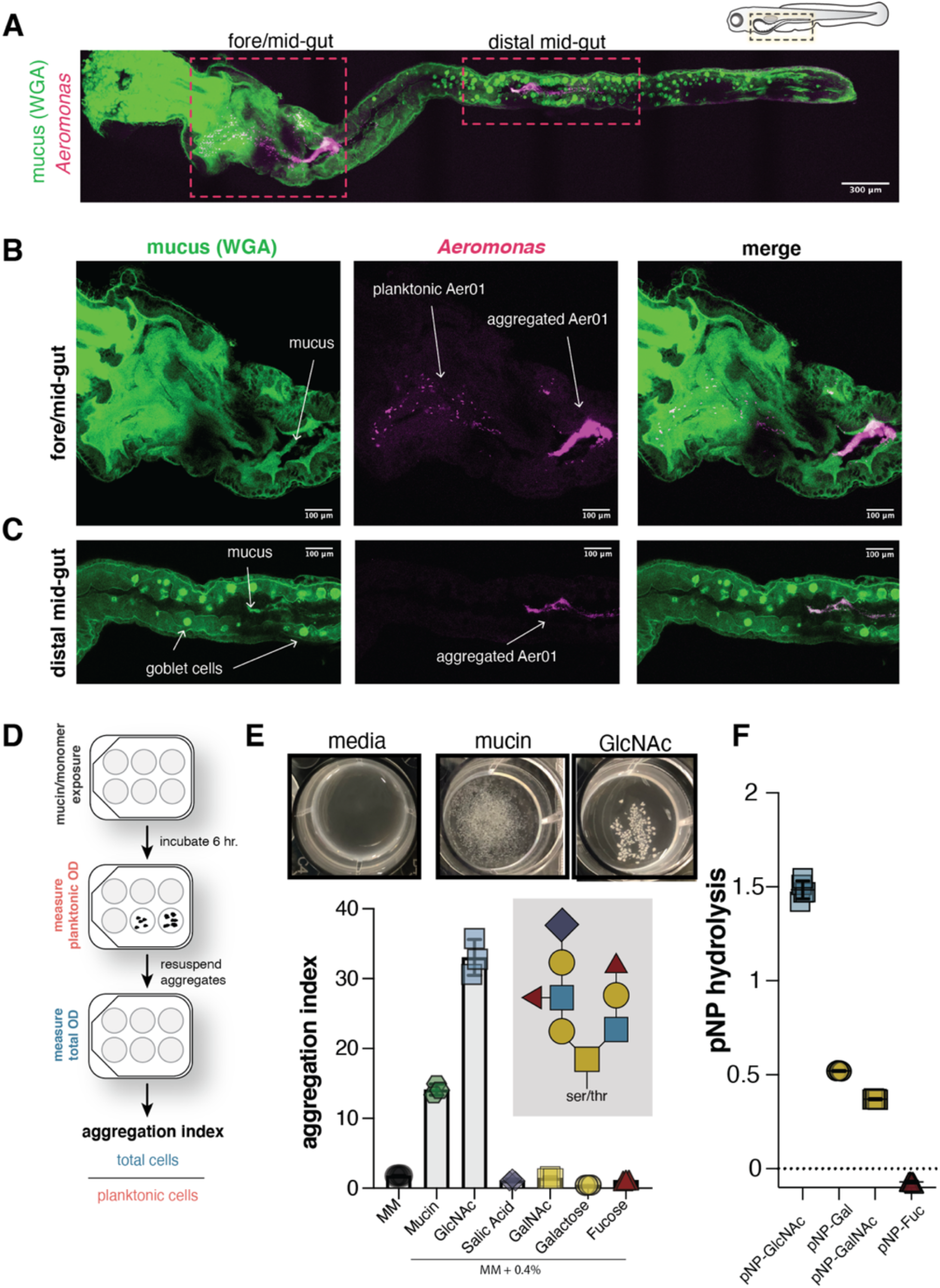
Aer01 associates with host intestinal mucus and aggregates in response to its mucolytic activity. **(A-C)** Confocal microscopy of fixed 5 dpf larval zebrafish intestine indicating Aer01(anti-dTom, magenta) and mucus (WGA-488, green) distribution (a) throughout the entire larval zebrafish gut, (b) the fore/mid- gut, and (c) distal mid-gut region. **(D)** Experimental design of culture-based aggregation assay and calculation of aggregation index. **(E)** Top: representative images of mucin- and GlcNAc-mediated Aer01 aggregation after 6 hr exposure. Bottom: mean aggregation index of Aer01 exposed to PGM or its monomeric glycan components (N=3). Insert: generic schematic of mucin O-linked glycan structure. **(F)** Aer01 cleave of mucin O-linkages was assayed by growing in the presence of 0.2% 4-Nitrophenyl (pNP) substrates for 6 hr. (N=5). Values presented represent the change in OD_405_ between the end and beginning of the experiment. The mean values are indicated.

Mucus accumulated in the lumen of the anterior and proximal end of the midgut (**Fig. 1B**, left). The majority of Aer01 cells were aggregated in the same proximal midgut region as mucus (**Fig. 1B**, right), with a small number of planktonic Aer01 present in the foregut (**Fig. 1B**, middle). This distribution of Aer01 in fixed tissue mirrors previous live imaging observations of Aer01 colonization dynamics (Schlomann et al., 2019; Wiles et al., 2016). The planktonic Aer01 may represent recently immigrated bacteria, as motility is critical for Aer01 immigration (Robinson et al., 2018, 2021). Additionally, we observed small Aer01 aggregates in the same region as mucus in the distal midgut lumen (**Fig. 1C**), which are likely mucus-encased Aer01 undergoing excretion.

These data indicate a strong spatial relationship between Aer01 aggregates and GlcNAc containing host mucus, and suggests mucus may promote Aer01 aggregation by passively encapsulating planktonic Aer01 or acting as a molecular signal that modulates Aer01 aggregation in the intestine.

### Mucin exposure in culture stimulates Aer01 aggregation

Since mucin is the major component of mucus, we next tested if mucin was sufficient to promote Aer01 aggregation in culture. We developed a culture-based approach outlined in **Fig 1D** to visualize and measure Aer01 aggregation upon exposure to minimal medium (MM) supplemented with commercially available porcine gastric mucin (mucin or PGM). In the presence of mucin, Aer01 forms dense, mucin-bound aggregates (**Fig. 1E**, top: mucin and **Fig. S1**) within 6 hours that are not formed in MM alone. Further, Aer01 aggregated in mucin concentrations as low as 0.02%, suggesting aggregation was not due to polymer-driven depletion aggregation in the viscous mucin medium (Azimi et al., 2021; Michaels et al., 2018) (data not shown).

To quantify the extent of aggregation under these conditions, the optical density of the planktonic fraction was measured, and then the bacterial aggregates were gently resuspended to allow a measurement of the total bacterial population density. A ratio of the cell densities (total/planktonic OD_600_) was calculated, which we report as the “aggregation index,” with increased aggregation corresponding to a higher aggregation index. Consistent with our visual assessment, we measured a higher aggregation index for Aer01 in mucin than in MM alone (**Fig. 1F**, bottom). Together, these data suggest Aer01 aggregation is an active response to mucin, suggesting this trait may play a role in Aer01’s spatial organization and distribution in the intestinal lumen.

### Mucolytic byproducts drive Aer01 aggregation in culture

To gain further mechanistic insights into Aer01’s response to mucin, we next asked if Aer01 encodes putative mucolytic enzymes involved in degrading glycans O-linked to mucin. Bioinformatic analysis indicates Aer01 encodes several glycoside hydrolase families (GH2, GH18, GH20, GH84, and GH129) employed by bacteria to hydrolyze intestinal mucin at GlcNAc, Gal, and GalNAc linkages (Mondal et al., 2014; Ndeh & Gilbert, 2018), suggesting Aer01 aggregation may be driven in part by sensing the activity of its own mucin glycan hydrolysis. Consistent with Aer01’s GH profile, Aer01 cleaved 4-Nitrophenyl (pNP) substrates containing GlcNAc, Gal, and GalNAc, but not Fuc (**Fig. 1F**), showing the highest hydrolytic activity against pNP-GlcNAc.

Because Aer01 can hydrolyze pNP substrates commonly used to assay mucin degrading capabilities, we next assayed Aer01 aggregation in MM containing 0.4% of each major O-linked glycan found on mucins: GlcNAc, Gal, GalNAc, Fuc, or SA. While several glycans supported Aer01 growth (albeit poorly due to the minimal medium growth conditions), GlcNAc alone was sufficient to drive Aer01 aggregation (**Fig. 1E**). These data suggest Aer01 aggregation in response to mucin may be driven by release of GlcNAc glycans produced Aer01’s GH activity.

### Experimental evolution reveals mucin-responses as a key determinant of Aer01 spatial organization in the intestine

To identify genetic pathways involved in Aer01 responses to mucin, we performed serial selections for Aer01 populations that remain planktonic in culture when exposed to GlcNAc, the key mucin glycan sensed by Aer01. For the serial selection, we exposed Aer01 to 0.4% GlcNAc as in the aggregation assay (**Fig. 1E**) and selected unaggregated Aer01 from the planktonic fraction after 6 hr (**Fig. 2A**, left). This planktonic sample was expanded by growing overnight in LB and again transferred back into GlcNAc medium for a total of 7 passages. Individual isolates were also selected from each passage by plating on LB agar. By the 5^th^ passage, no Aer01 aggregates were observed in GlcNAc or mucin (**Fig. 2A**, right). Thus, we named the non-responsive isolates from these populations “<u>m</u>ucin-<u>b</u>lind” (MB).

**Figure 2.**
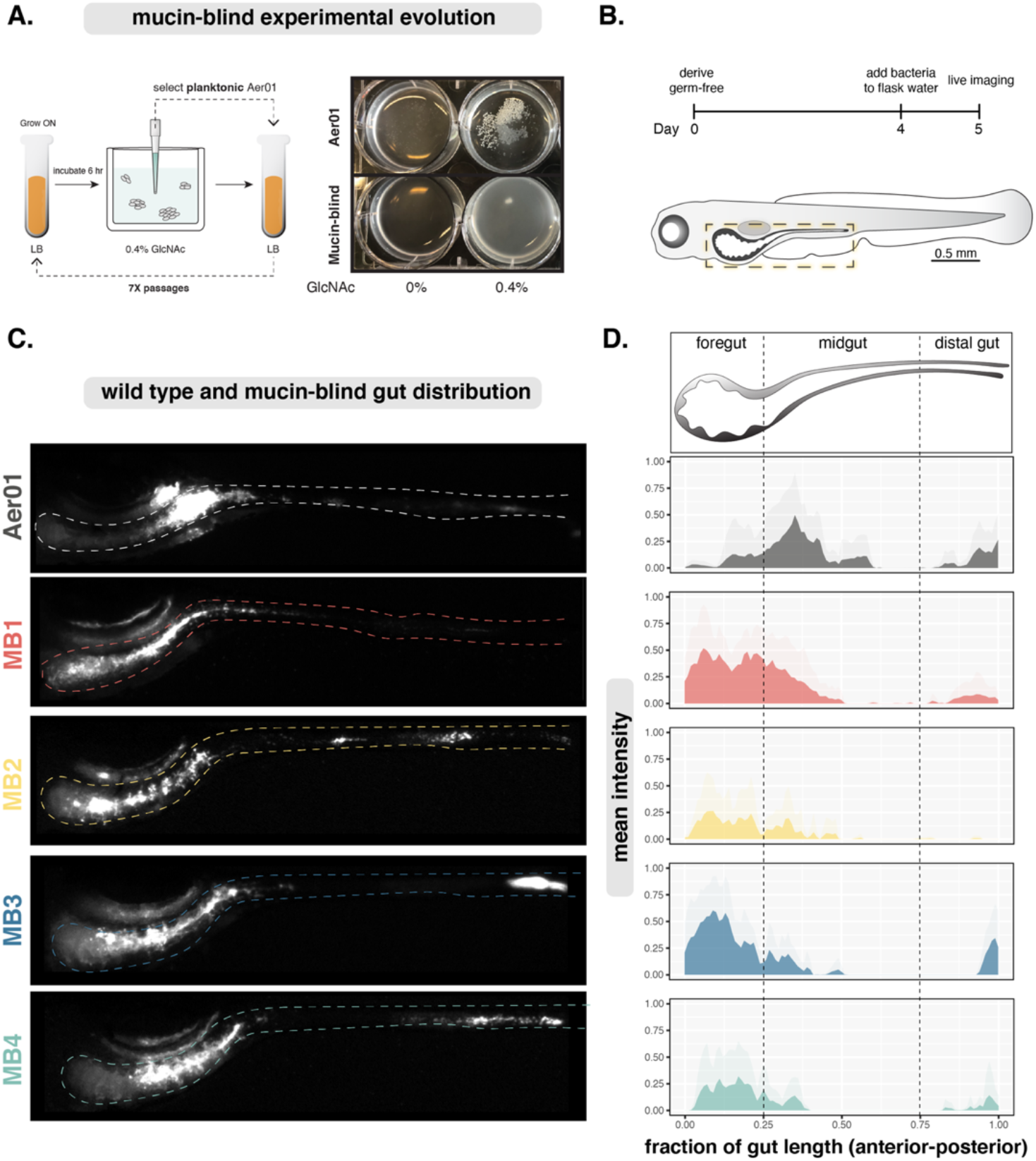
Defects in mucin responses alter Aer01 spatial organization in the intestine. **(A)** Left: Experimental evolution schematic; Aer01 was exposed to minimal media (MM) supplemented with 0.4% GlcNAc for 6 hr and planktonic cells were selected, regrown overnight in LB, then repassaged in MM-GlcNAc for a total of 6 passages. Evolved isolates from this experiment are designated mucin blind (MB). Population and isolate samples from each passage were cryopreserved. Right: representative images of Aer01 aggregates and mucin-blind passage 5 isolate collection 6 hours after GlaNAc exposure. **(B)** Time-line for WT and MB mono-association experiments to compare population structures in the intestine. **(C)** Representative live stereoscope microscopy images of larval zebrafish mono-associated with wild type or indicated MB isolates 24 hpi (5 dpf). The approximate intestinal boundary is outlined and corresponds the region measured for distribution analysis in (d). **(D)** Quantification of biogeography from lives images in (c). Mean pixel intensities (microbial abundance) are plotted along the normalized gut length and standard deviation at each point plotted in the corresponding lighter color. Measurements were conducted in Fiji. Aer01 (n=7), MB1 (n=7), MB2 (n=7), MB3 (n=5), MB4 (n=7).

Next, we tested if the genetic pathways disrupted in the culture-based experimental evolution screen contributed to Aer01’s aggregation and population structure in the intestine. We randomly selected four MB isolates, two from passage 5 (MB1 and MB3) and two from passage 6 (MB2 and MB4), and assayed their spatial organization throughout the entire larval zebrafish gut using live fluorescence stereomicroscopy. Each MB isolate was labeled with dTom for live imaging using molecular genetic tools developed by our lab (Wiles et al., 2018). MB isolates were monoassociated with germ free larval zebrafish at four dpf and imaged 24 hours later (**Fig. 2B**). The mean normalized pixel intensity corresponding to microbial abundance was plotted across the normalized gut length along with the standard deviation at each position for WT and our MB isolates of Aer01. As expected, WT Aer01 formed large aggregates in the anterior midgut region with few planktonic cells in the bulb (**Fig. 2C** and **D**; Aer01). Strikingly, all four MB isolates exhibited altered distributions with an anterior displacement into the foregut region (**Fig. 2C** and **D;** MB1, MB2, MB3, MB4). These data suggest that Aer01’s ability to sense and respond to mucin or mucin-associated glycans shapes its spatial distribution and organization in the gut.

### MB mutations converge on a secondary messenger-regulated adhesin

To uncover the genetic mutations associated with our four MB isolates, we sequenced and compared their genomes to the parental strain using *breseq* (Deatherage & Barrick, 2014). In total, four single nucleotide polymorphisms (SNPs) and one indel were identified. The locus ID, putative functions, and mutations for each isolate are shown in **Fig. 3A**. We found the genomic changes were in genes that function in aggregation/biofilm formation or c-di-GMP (cdG) metabolism, a key regulator of aggregation.

**Figure 3.**
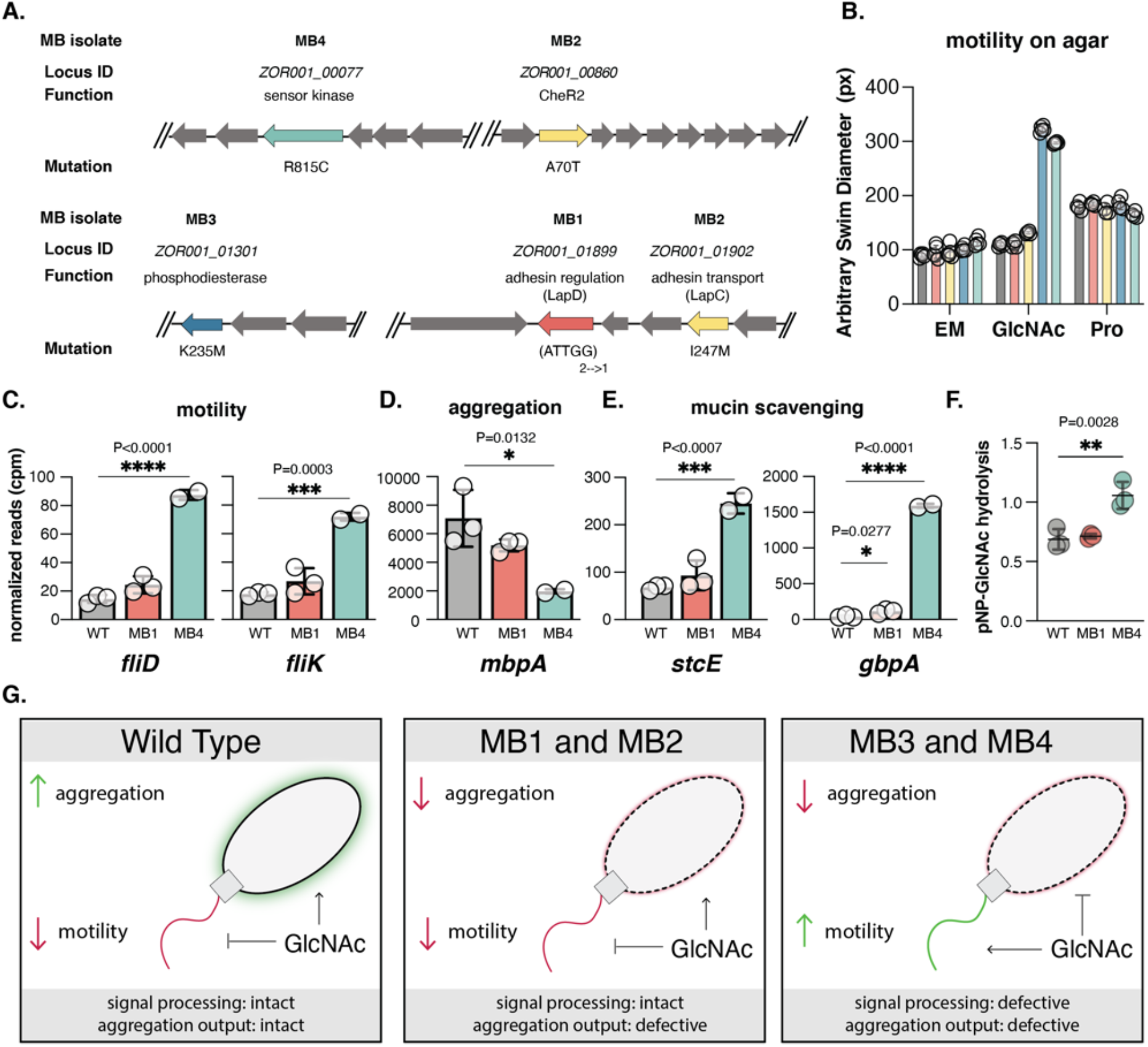
Genomic analysis of experimentally evolved isolates reveals molecular basis for mucin-mediated aggregation defects. **(A)** Chromosomal organization of mutations in select MB isolates were identified using *breseq*. Mutations are colored by isolate (red: MB1; yellow: MB2; blue: MB3; teal: MB4) and the locus ID, putative function, nucleotide substation, and corresponding amino acid change listed when applicable. **(B)** Swim diameter of WT and MB isolates in MM alone (N=5 replicates) or supplemented with 0.4% GlcNAc (N= 5 replicates) or 1 mM proline (N=4 replicates) after 20 hr incubation at 30°C. (**C-E**) RNAseq results of indicated genes from WT (N=3), MB1 (N=3), and MB4 (N=2) incubated in 0.4% GlcNAc MM for 6 hours. Mean and standard deviations are indicated. Ordinary one-way ANOVA with multiple comparisons. **(F)** Hydrolytic activity of WT, MB1, and MB4 against the pNP-GlcNAc substrate shows MB4 exhibits increased pNP hydrolysis, supporting RNA-seq analysis (N=3). **(G)** Graphical model of MB mutant classes based on in vitro aggregation and motility analysis. MB isolates contained mutations linked to defects in a c-di-GMP-regulated adhesin output pathway (MB1 and MB2) or signal processing pathways involved in c-di-GMP metabolism (MB3 and MB4). For MB1 and MB2, signal processing of GlcNAc remains intact, but defects in localization or secretion of the adhesin MbpA disrupt aggregation. For MB3 and MB4, dysregulation of GlcNAc-mediated c-di-GMP signaling leads to increased motility and altered gene regulation.

MB1 and MB2 harbor mutations in a widely distributed bacterial biofilm regulatory pathway called the Lap system that was first described in *Pseudomonas fluorescens* (Collins et al., 2020). In this system, a giant adhesin (LapA) is transported to the cell surface via a cognate Type 1 Secretion System (T1SS) secretion apparatus (LapBCE). Cell surface levels of LapA are controlled by the activity of the cdG-sensing regulatory protein LapD that represses the activity of a LapA-targeting protease, LapG. Cleavage of LapA by LapG releases LapA from the cell surface and inhibits LapA function in biofilm formation. MB1 contains a 5 nucleotide deletion in *lapD* at nucleotides 76-80 of the 1926 nucleotide gene. This indel generates an early frameshift predicted to generate a loss of function and suggesting MB1 is unable to sustain normal adhesin levels due to constitutive activity of the adhesin-targeting protease. As in other Lap systems, the putative LapG target, *ZOR001_01898*, is encoded adjacent to *lapD* and the other Lap components.

The MB2 SNP generates a I247M mutation in the LapC membrane fusion protein (MFP) component of the T1SS apparatus. In other MFPs, mutations in this region can diminish protein secretion by disrupting interactions between the MFP and its target outer membrane protein (Kim et al., 2016), suggesting this mutation may block transport or inhibit retention of the adhesin. MB2 also contains a mutation in CheR2, but since MB2 phenocopies MB1, this mutation likely does not contribute to the MB2 phenotype. We thus named the Aer01 genes at this locus identified through our <u>m</u>ucin-<u>b</u>lind screen as MbpABCDEG, following the convention of the Lap component letter designations.

MB3 and MB4 contain mutations in predicted cdG metabolizing and sensory signaling pathways. MB3 has a K235M mutation in the HD-related output domain (HDOD) of ZOR001_01301, a predicted phosphodiesterase (PDE) enzyme that hydrolyzes cdG. This protein has 64% amino acid similarity to CdgJ, a PDE involved in *Vibrio cholerae* biofilm formation (Jones et al., 2015). HDODs are often associated with PDE and DGC enzymes, but their role in modulating cdG metabolism is poorly understood.

MB4 has a R815C mutation in the dimerization and phosphor-acceptor domain of an atypical two-component system (TCS) histidine kinase (HK), ZOR0001_00077. TCSs are regulatory phosphor-relay hubs comprised of an environment-sensing HK and a cognate response regulator (RR). Most TCS HK-RR pairs are typically co-translated. In the case of ZOR0001_00077, the co-transcribed RR is a predicted cdG-degrading PDE, suggesting this HK is involved in relaying environmental information such as GlcNAc availability to modulate Aer01 cdG metabolism and thus Aer01 biofilm formation or motility. We thus named this sensor kinase “<u>m</u>ucin <u>b</u>lind <u>k</u>inase”, MbkS. Together, genetic analysis of our MB isolates suggests the model that in wild-type Aer01 the TCS HK (MbkS) and PDE (ZOR001_01301) respond to environmental GlcNAc, leading to increased cell surface levels of MbpA via MbpD.

### Aer01 uses distinct molecular signaling pathways for adapting to extra-host and host-associated lifestyles

Bacteria transition from the planktonic to aggregated or biofilm lifestyle by decreasing their motility and increasing levels of cell surface-associated adhesins. The secondary messenger cdG is a key regulatory molecule in this process, with cdG synthesis driving aggregation and depletion stimulating motility. Genomic analysis of our MB isolates identified two isolates (MB3 and MB4) with mutations in sensory domaincontaining proteins linked to cdG metabolism and two isolates with defects in a cdG-sensing aggregation pathway downstream of GlcNA sensing (MB1 and MB2). MB1 and MB2 mutations in aggregation output suggest these isolates sense and relay environmental GlcNAc availability similarly to WT, but cannot deploy MbpA in response to this cue. Conversely, the MB3 and MB4 mutations alter proteins with sensory domains putatively involved in sensing environmental GlcNAc. Thus, we predicted MB1 and MB2 decrease their motility in GlcNAc like WT, but fail to aggregate, while MB3 and MB4 remain motile and never initiate transition from the planktonic to aggregated lifestyle. In support of this idea, swim plate analysis on minimal medium or minimal media supplemented with 0.4% GlcNAc indicate MB3 and MB4, but not MB1 or MB2, are motile in the presence of GlcNAc (**Fig. 3B**; GlcNAc). All MB isolates swim similarly to WT in the presence of proline, a known stimulant of Aer01 motility (Robinson et al., 2021), suggesting MB3 and MB4 signaling defects are specific to GlcNAc.

To further explore Aer01 responses to GlcNAc, we performed RNA-seq analysis of WT, MB1, and MB4 grown in MM supplemented with GlcNAc. Consistent with our swim plate assays, RNA-seq analysis of WT, MB1, and MB4 transcripts following GlcNAc exposure indicated MB4 expresses motility-associated genes such as *fliD* and *fliK* at higher levels than WT or MB1, which are not significantly different (**Fig. 3C**). Conversely, we also found MB4 exhibits defects in expression of aggregation associated genes such as *mbpA* compared to WT and MB1 (**Fig. 3D**). In support of this data, we observed levels of *mbpA* promoter activity in WT, MB1, and MB4 Aer01 carrying a P*_mbpA_::mNeonGreen* transcriptional reporter that mirrored our RNA-seq analysis (**Fig. S2**). Importantly, although MB1 expresses *mbpA* similarly to WT, loss of *mpbD* due to an indel inhibts MbpA function. This defect can be rescued by expressing the wild-type *mbpD* allele in the MB1 background (**Fig. S3**). Thus, although MB1 and MB4 exhibit differences in GlcNAc-mediated motility in our plate-based assay, both isolates have disrupted MbpA activity. Additionally, general cdG signaling defects are not sufficient to inhibit GlcNAc-mediated aggregation as a hyper-motile Aer01 *ΔspdE* mutant (Robinson et al., 2021) aggregates like wild type upon GlcNAc exposure (**Fig. S4**). These data highlight the roles of distinct microbial sensory pathways in tuning the behavior of a beneficial bacterium in response to chemical cues of host proximity (proline) versus host-association (mucin and GlcNAC).

### GlcNAc signal processing defects stimulate mucin degradation pathways

RNA-seq analysis revealed that MB4 exhibits marked up regulation of genes involved in mucin degradation and GlcNAc scavenging. Both *gbpA* and *stcE* are upregulated in MB4 compared to WT cultured in GlcNAc supplemented minimal medium (**Fig. 3E**). GbpA is a copper-dependent lytic polysaccharide monooxygenases (LPMO) first identified as a GlcNAc-binding protein from *Vibrio cholerae* that recognizes GlcNAc moieties on mucin and has since been shown to degrade GlcNAc-containing polymers such as chitin (Bhowmick et al., 2008; Loose et al., 2014) and stimulate intestinal epithelial proliferation in the gnotobiotic zebrafish intestine (Banse et al., 2022).

Like *gbpA* from *Vibrio*, Aer01 *gbpA* contains a Cyclic di-GMP-I riboswitch (RF01051) in its putative 5’-UTR, which, in *Vibrio* increases translation of *gbpA* when cdG levels are low (Sudarsan et al., 2008). StcE is a mucin-selective metalloprotease from *Escherichia coli* that cleaves mucins as well as O-glycosylated cell surface glycoproteins (Grys et al., 2006; Hews et al., 2017; Malaker et al., 2019). Consistent with the idea that MB4 exhibits increased mucolytic potential, it cleaved GlcNAc pNP substrates more efficiently than WT (**Fig 3F**). Together, these data suggest MbkS typically tunes Aer01 cdG levels in response to GlcNAc availability in the intestine, adapting Aer01’s physiological strategies to take advantage of available GlcNAc polymers through transcriptional regulation of mucin-degrading secreted factors. Collectively our genomic analysis and culture-based assays suggest MB isolates exhibit defects in sensing (MB3 and MB4) or responding (MB1 and MB2) to GlcNAc (**Fig. 3G**).

### MbpA function drives aggregate formation in the intestine

At the population level, MB isolates exhibit altered spatial organization, being displaced from their typical location in the proximal midgut into the foregut (**Fig. 2C**). We next sought to explore if mucus sensing and aggregation are important for bacterial behaviors in the mucus-rich intestine. We characterized the intestinal distributions and colonization dynamics of MB4 and MB1. In our culture-based assays both MB isolates fail to aggregate when exposed to mucin or GlcNAc. Notably, in these assays MB4 is hyper motile in the presence of GlcNAc whereas MB1 motility resembles WT. Using fluorescence confocal microscopy on fixed intestines as previously (**Fig. 1A**), we found MB1 and MB4 were largely anteriorly displaced into the foregut (**Fig. 4A)** but co-localized to some extent with mucin in the midgut (**Fig. 4 B and C**). Strikingly, both MB isolates appeared to be more planktonic and form smaller aggregates than their wild type counterpart.

**Figure 4.**
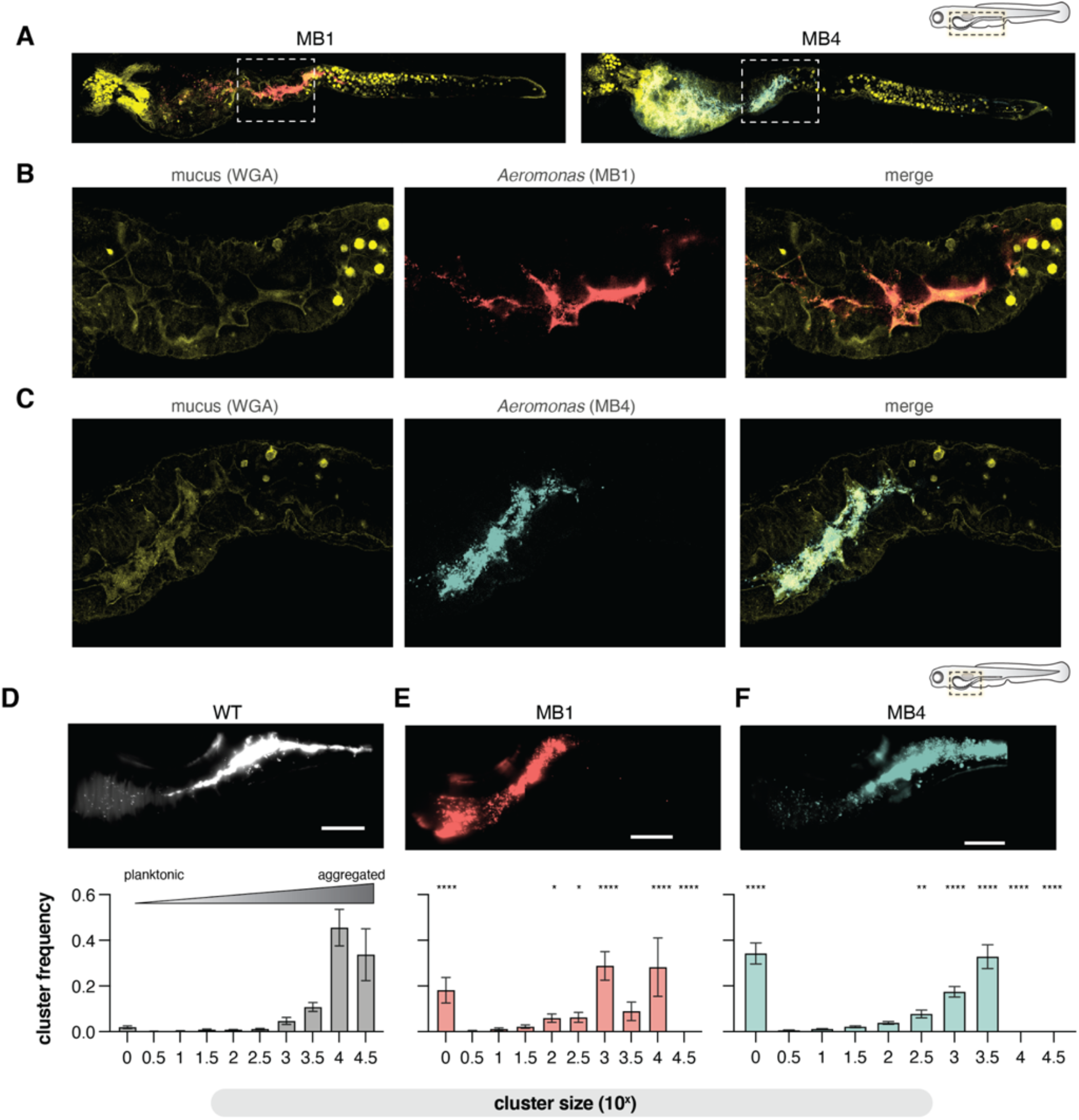
MB isolates are more planktonic in the mucus-rich lumen. **(A)** Confocal microscopy of fixed 5 dpf larval zebrafish intestine indicating MB1 (anti-dTom, red), MB4 (anti-dTom, blue), and mucus (WGA-488, yellow) distribution throughout the entire larval zebrafish gut. **(B-C)** zoom in of the fore/mid-gut indicated in A and B (boxed regions) suggest decreased aggregation alters MB1 and MB4 population structures. **(D-F)** Top: maximum intensity projections from live light sheet fluorescence microscopy of the distal foregut to proximal mid gut region of the larval zebrafish intestine showing the typical population structures of WT, MB1, and MB4. Bottom: The probability of being in an n-cell cluster for WT, MB1, and MB4. The mean and standard deviation are indicated (WT, N=11; MB1, N=7, MB4, N=6). Two-way ANOVA with multiple comparisons to WT (*p <0.05, **p<0.01, ***p<0.0001). Scale bars=100 μm.

Light sheet fluorescence microscopy revealed striking differences between MB1 and MB4 behaviors in the intestinal foregut environment. Consistent with our model that MB1 can sense mucus and respond by downregulating motility, but is defective in mucus-induced aggregation (**Fig. 3G**), MB1 was found predominately in small aggregates and as single non-motile planktonic cells (**Movie S3**). Conversely, MB4 was largely motile, with planktonic cells frequently swimming into the epithelium (**Movie S4**).

To quantify these population structures, we used live light sheet fluorescence microscopy to image a 800 μm region of interest corresponding to the anterior bulb and distal midgut of larval zebrafish monoassociated with WT, MB1 or MB4. The three-dimensional volume of the intestinal lumen was analyzed and Aer01 cells identified and categorized using custom software described in Sundarraman et al. 2022. Here we compare the population structure of MB1 to our previously reported distributions of WT and MB4. Representative maximum intensity projections along this area are shown in **Fig. 4 D-F**. We found WT Aer01 are most often assembled into large multi-cellular aggregates of 10^4^ and 10^4.5^ cells (**Fig. 4D**), with planktonic cells (10^0^) rarely observed. The cluster size distribution is shifted in MB1 (**Fig. 4E**) and MB4 (**Fig. 4F**). MB1 forms smaller (10^3^-10^4^ cell) clusters with non-motile planktonic cells (10^0^) dispersed throughout the intestine. The motile MB4 is predominantly planktonic (10^0^) with some smaller cell clusters (up to about 10^3.5^ cells), consistent with our confocal imaging. Together, these data highlight the crucial role of a microbial sensory pathway in determining symbiont intestinal aggregation and population structure along the length of the intestine.

### Mucin regulatory pathways tune symbiont inflammation potential independently of motility

We had previously shown that bacterial motility is necessary and sufficient for a pro-inflammatory *Vibrio* species to elicit intestinal inflammation in the zebrafish intestine (Wiles et al., 2020). We thus anticipated that the highly motile MB4 would elicit more intestinal inflammation than its non-motile WT or MB1 counterparts. To test this hypothesis, we enumerated intestinal neutrophil abundance using transgenic zebrafish *Tg(mpx:gfp*) with GFP labeled neutrophils after mono-association with either WT, MB1, or MB4 (**Fig. 5A,B**). Surprisingly, we found both MB1 and MB4 increase intestinal inflammation compared to WT (**Fig. 5C**) despite their clear motility differences in the host, suggesting motility is not necessary for Aer01-induced inflammation. The increase in intestinal inflammation was apparent when neutrophils were quantified along the entire length of the intestine, as shown, or if neutrophil enumeration was restricted to the mid and distal intestine (data not shown). For subsequent experiments, neutrophil quantification was restricted to the mid and distal intestine. We found no differences in intestinal colonization between WT, MB1, or MB4 (**Fig. 5D**), indicating heightened inflammation was not due to MB1 or MB4 intestinal overgrowth but rather to alterations in symbiont-innate immune system interactions.

**Figure 5.**
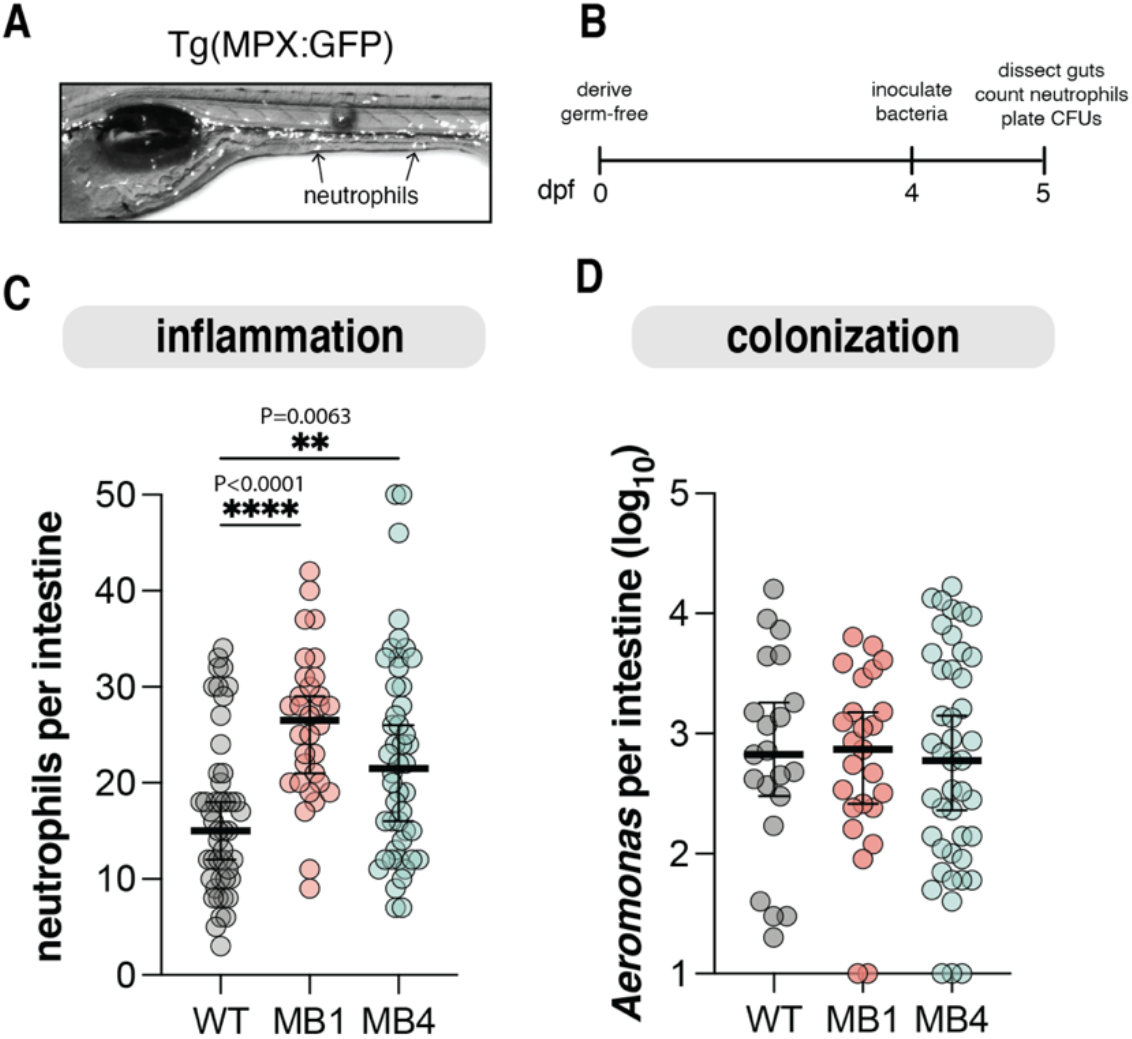
Mucin-blind isolates are more proinflammatory. **(A**) Representative live fluoresence stereoscope microscopy image of Tg(MPX:GFP) larval zebrafish 5 dpf. **(B)** Schedule of GF derivation and Aer01 mono-association for Aer01 abundance and host intestinal inflammation assays. **(C)** Neutrophil abundance from intestines of larval zebrafish mono-associated with the indicated Aer01 strain. Line and error bars represent median with 95% CI. For all graphs each dot represents one fish; n ≥ 32 from at least 3 independent experiments. (WT, N=47; MB1, N=32; MB4, N=48). Ordinary one-way ANOVA was performed for *Aeromonas* abundance and neutrophils (P values: ****>0.0001, **=0.0063). **(D)** Abundance of WT, MB1, and MB4 24 hr following mono-association as determined by plating a sample from dissected, homogenized intestines on TSA agar plates and calculating CFUs. Line and error bars represent median with 95% CI. WT, N=23; MB1, N=26; MB4, N=46.

### MbpA regulation limits Aer01’s inflammation character

Our unexpected observation that non-motile MB1 caused similar inflammation to the hyper-motile MB4 isolate motivated us to further explore the idea that Aer01’s inflammatory character is limited largely by proper MbpA regulation inside the gut environment. To accomplish this, we leveraged previous studies (Boyd et al., 2012; Newell et al., 2011) on LapD-LapG regulation of LapA (MbpD-MbpG and MbpA in Aer01) to design a synthetic construct that can convert WT Aer01 to an MB1 phenotype upon addition of the inducer anhydrotetracycline (aTc). In the Lap system, plasmid-based over-expression of the LapG protease can overcome LapD inhibition of LapG proteolysis, which phenocopies a genetic loss of *lapD*. MB1 contains a LOF in *mbpD* allowing constitutive MbpG activity (**Fig. 6A**; left, MB1). Thus, we reasoned that induced expression of *mbpG* in WT Aer01 would result in loss of MbpA function and Aer01 aggregation similarly to the MB1 isolate (**Fig. 6A**; right, WT^mbpG^). A schematic representation of this inducible system is shown in **Fig 6B**: addition of the inducer aTc drives mbpG and mNeonGreen co-expression (mbpG) or only mNeonGreen as a control (empty). These constructs also encode constitutively expressed dTom for tracking live Aer01 cells in the host. Consistent with our model outlined in **Fig. 6A**, *mbpG* induction inhibited aggregation of Aer01 in the presence of GlcNAc (**Fig. 6C**, top), phenocopying MB1. Additionally, inducing *mbpG* expression following Aer01 aggregation dispersed the preformed aggregates (**Fig. 6C**, bottom), revealing that MbpA regulation is also involved in aggregate dispersion.

**Figure 6.**
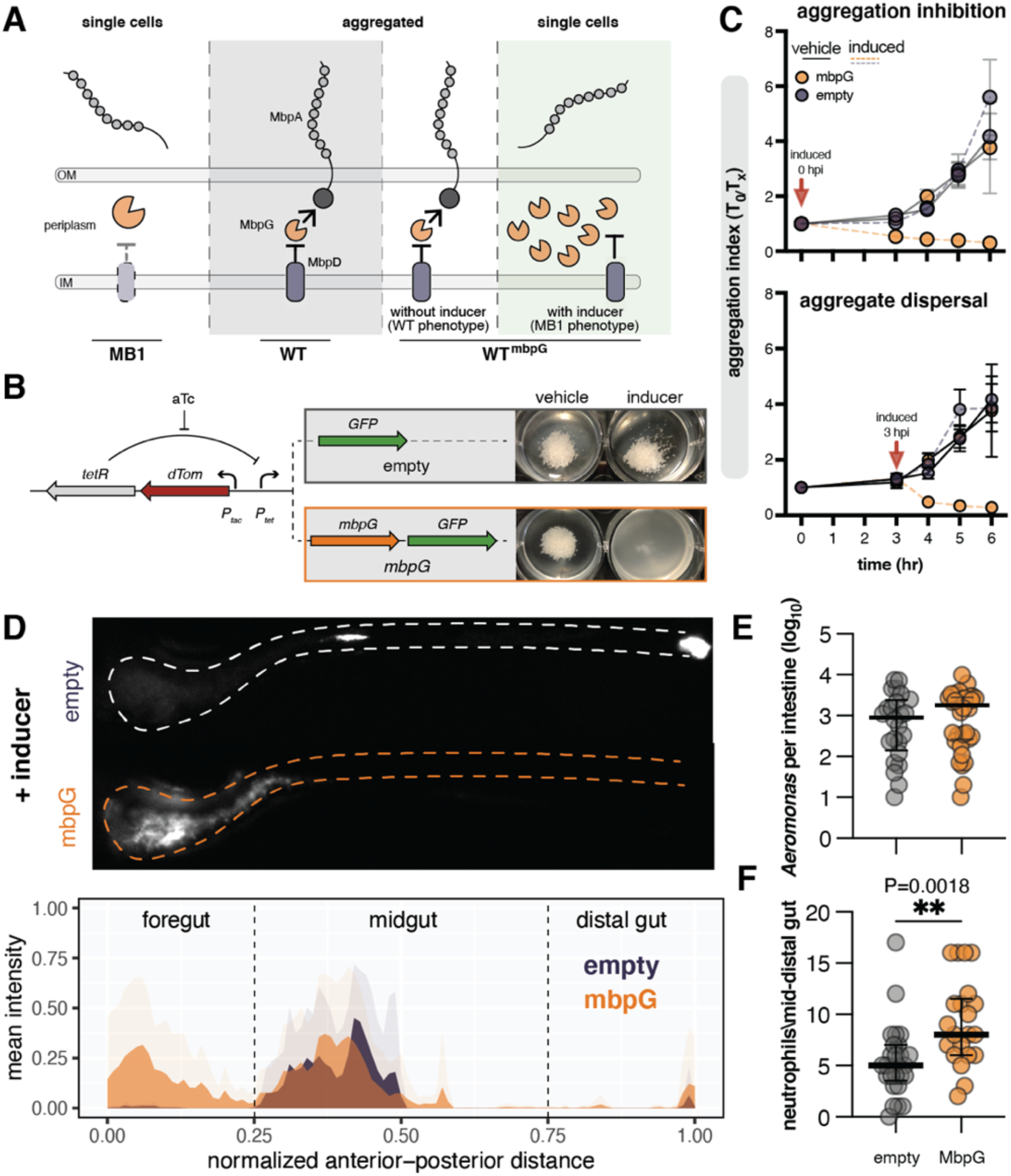
Disruption of MbpA function in vivo disrupts Aer01 biogeography and increases intestinal inflammation. **(A)** Schematic model of MbpA regulation. Middle (WT) Left (MB1): loss of the protease regulator MbpD in the MB1 mutant leads to constitutive protease activity and MbpA inactivation. induction cassette to disrupt MbpA function. Over expression of the MbpA-targeting protease MbpG. **(B)** Left: schematic representation of the engineered construct to inactivate MbpA activity in WT Aer01. The addition of aTc derepressed the expression of GFP control (empty) or *mbpG* and GFP (mbpG). Right: induction of the *mbpG*, which cleaves and inactivates MbpA, is sufficient to suppress GlcNAc-mediated WT Aer01 aggregation in culture. **(C)** Induction of *mbpG* inhibits aggregation (top) and disperses formed aggregates (bottom). Vehicle (MM, solid lines) or aTc (aTc in MM, dashed lines) at the time of MM-GlcNAc exposure (top) or after 3 hr. after exposure when Aer01 formed aggregates (bottom). **(D)** Inhibition of MbpA activity via *mbpG* induction alters WT Aer01 population structure in the intestine, leading to displacement into the foregut region. Top: representative image WT carrying either the empty or mbpG detailed in (B). Bottom: quantification of WT population structure across the entire gut. Solid colors indicate the mean value at the normalized distance with transparent colors indicating the standard deviation at that position. WT^empty^ N=5; WT^mbpG^, N=11. **(E)** Intestinal abundance of Aer01 carrying either the empty or mbpG constructs. Line and error bars represent median with 95% CI. Unpaired t test. WT^empty^ N=24; WT^mbpG^, N=34. **(F)** Inactivation of MbpA function increases the inflammation character of WT Aer01. Intestinal inflammation is indicated by neutrophil abundance in the mid to distal gut. Line and error bars represent median with 95% CI. WT^empty^ N=25; WT^mbpG^, N=21.

Next, to test the functionality of our synthetic construct inside the host, we assayed Aer01 distribution, colonization, and host inflammation as previously by mono-associating larval zebrafish 4 dpf with WT Aer01 carrying either the *mbpG* or empty synthetic cassette and inducer. We found *mpbG* induction for 24 hours caused similar anterior displacement into the intestinal bulb region as seen in the MB1 isolate (**Fig. 6D**). The empty cassette control strain formed mid-gut aggregates similar to those previously observed with WT Aer01 (**Fig. 6D**). As expected, *mbpG* induction did not impact colonization levels (**Fig. 6E**). Phenocopying MB1, *mbpG* induction led to increased intestinal neutrophil abundance in the mid and distal gut (**Fig. 6F**). Together, these data indicate that symbiont adhesin regulation in response to host-associated environmental cues limits the inflammatory character of Aer01.

### A human gut symbiont MbpA analog reduces MB1 induced gut inflammation

SMART analysis (Letunic et al., 2021) of MbpA’s domain architecture revealed it contains 20 repeats of a putative glycan-binding parallel beta-helix (PbH1) domain, 19 of which are identical in sequence (**Fig. 7A**, MbpA). To determine if other host-associated microbes encode putative adhesins with similar architecture, we again utilized the SMART database to identify likely adhesins containing PbH1 domains. We focused on adhesins belonging to the Type 5a Secretion System (T5aSS) in Gram-negative bacteria because adhesins of this group are easily identified by the presence of a C-terminal autotransporter domain. Our search for proteins with a C-terminal Autotransporter domain and at least 1 PbH1 domain returned >2,700 adhesins from various Gram-negative bacteria strains that ranged from 559 to 16,945 amino acids in length. Approximately 75% of the bacterial species encoding these adhesins have been identified in human ileum, colon, or stool samples via 16S sequencing according to the IBD TaMMA database (Massimino et al., 2021). We provide a table with the uniport code, species, putative function, and amino acid sequences of MbpA-like adhesins identified (**Table S1**).

**Figure 7.**
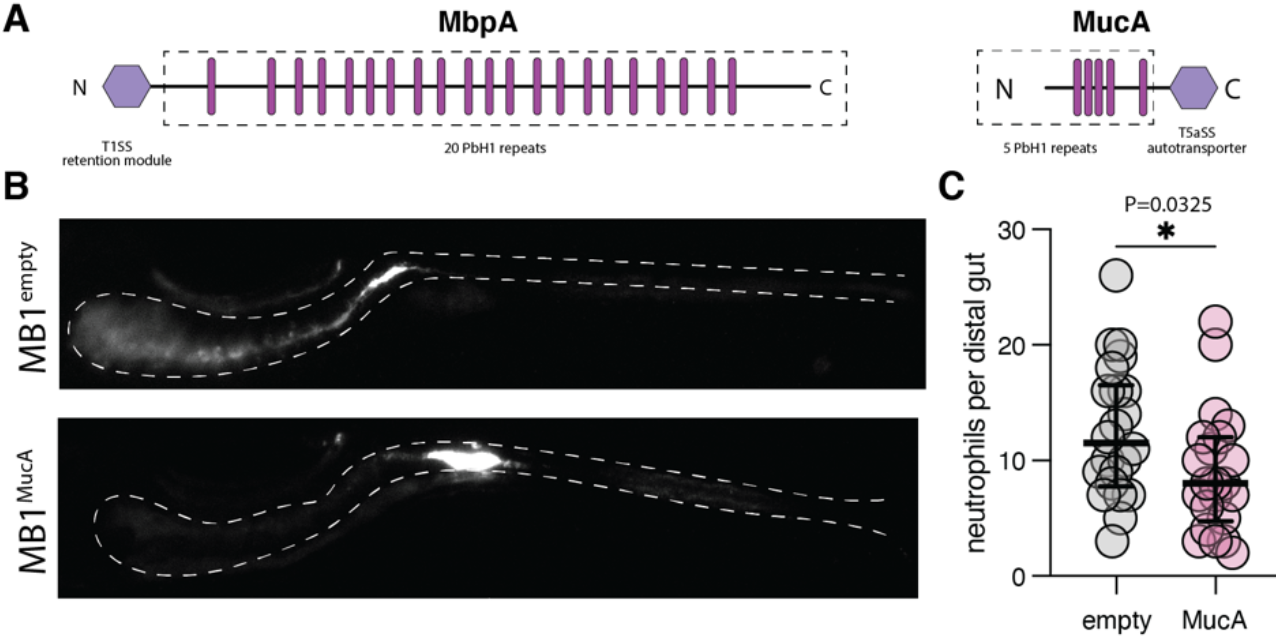
Expression of the *Akkermansia* adhesin MucA rescues MB1 population structure and reduces MB1 inflammation. **(A)** The PbH1-containing adhesin MbpA and MucA contain similar domain architectures but are secreted and regulated via distinct mechanisms (T1SS and T5aSS). The dashed boxed region indicates putative surface exposed domains, the pink bars approximate the location of each PbH1 domain as indicated by SMART analysis, and purple hexagons are the domains involved in retaining the respective adhesins at the cell surface. The size of each adhesin is scaled according to the amino acid length. **(B)** Expression of MucA rescues population structure defects associated with MB1 in the intestine as indicated by live fluorescence stereoscopy. **(D)** The *Akkermansia* adhesin MucA reduces intestinal inflammation associated with the MB1 evolved isolate. Neutrophil abundance was compared using an unpaired t test (N=22, each). Line and error bars represent median with 95% CI.

We hypothesized adhesins of this group likely function in microbial aggregation or binding glycan-rich substrates such as mucus. To test this idea, we sought to determine if an uncharacterized MbpA analog from *Akkermansia muciniphilia--Amuc_1620* (**Fig. 7A**, MucA for <u>muc</u>iniphilia <u>a</u>dhesin)—could rebalance MB1 population structure and inflammatory potential. Previous proteomic analysis indicated MucA is ~20 fold more abundant in the outer membrane of *Akkermansia* grown in mucin-rich growth conditions (Ottman et al., 2016), suggesting *Akkermansia* and Aer01 deploy functionally analogous adhesins to their cell surface upon mucin exposure. We reasoned Aer01 can display MucA at its cell surface because T5aSS adhesins are broadly encoded by Gram-negative bacteria (including *Aeromonas* species) and their transport and insertion into the outer membrane rely on the highly conserved Sec and Bam machinery. Importantly, unlike MbpA, MucA is not a substrate for the Lap system and its function is regulated independently of MbpG activity. This feature allows us to express MucA in the MB1 background (where MbpG is constitutively active) without impacting MucA function.

We used a similar cloning and induction strategy as detailed previously (**Fig. 6B**) to express the *mucA* open reading frame from *A. muciniphilia* ATCC BAA-835 under regulation of the tet operator, such that MucA protein would be induced upon addition of aTc. In support of the idea that *E. coli* and *Aeromonas* can translocate and display MucA, we observed the lab strain of E. coli (S17) carrying the *mbpA* construct formed filament-like aggregate structures upon *mbpA* induction (data not shown), further suggesting a role for MucA and MbpA-like adhesins in aggregation.

We next tested whether expression of the MbpA analog from *Akkermansia* in MB1 could repair the altered intestinal population structure and pro-inflammatory properties associated with the MB1 isolate. When induced in MB1 colonizing the zebrafish intestine, MucA largely restored MB1’s population organization and distribution, reducing the population of planktonic cells in the foregut and leading to the formation of large aggregates in the anterior midgut region (**Fig. 7C**), characteristic of the parental Aer01 (**Fig. 2C**). Concomitant with this rescue of population structure, MucA expression in MB1 significantly reduced the capacity of this strain to elicit intestinal inflammation. Together, these data suggest a broad role for MbpA-like adhesins in bacterial aggregation and host immunomodulation in response to mucins.

## Discussion

Mucin’s physical barrier function has long been appreciated for its ability to modulate the structure and function of microbiota (Hansson, 2012). In this study, we demonstrate the active role bacterial play in determining their population structure and host interactions within the gut. We show that the zebrafish mutualist Aer01 governs its gut distribution and inflammatory character in response to host mucin signals by regulating the activity of a large ~500 kDa adhesin we named MbpA. Microbial adhesins are well know for their role in biofilm formation and aggregation in culture, but their contribution to the organization and distribution of microbial communities in the intestine and corresponding impacts on host health are poorly understood. Our work demonstrates that Aer01 detects molecular cues from the healthy gut mucus ecosystem (GlcNAc) using a TCS sensor kinase (MbkS) to control various aspects of its physiology inside the host including tuning the transcription and cell surface localization of MbpA--an adhesin required for aggregation in response to GlcNAc.

Bacterial sensing of environmental cues such as mucin glycans through TCSs can rapidly alter transcription and physiological behaviors to change their interactions with the host immune system. For example, the TCS RprXY of Enterotoxigenic *Bacteroides fragilis* represses production of the metalloprotease *B. fragilis* toxin (BFT) in response to yet to be determined mucus signals, and induces BTF expression when the mucus ecosystem is compromised (Hecht et al., 2017). Similarly, in *Pseudomonas aeruginosa* mucin glycans act through the sensor kinase RetS to inhibit GacS and Type-6 secretion system (T6SS) activity and toxin secretion (Wang et al., 2020). RNA-seq analysis of WT Aer01 and the MB4 isolate exposed to GlcNAc in culture indicate MbkS activity modulates the transcription of *mbpA*, motility-associated genes (ie, *fliD* and *fliE*), and several mucin-degrading enzyme genes (*stcE* and *gbpA*), the latter of which induces proliferative responses in the zebrafish intestinal epithelium (Banse et al., 2022). In the gut, the motile MB4 isolate resists formation of large aggregates characteristic of its WT counterpart and elicits more intestinal inflammation. We initially hypothesized that MB4’s increased motility would render it especially pro-inflammatory, possibly through elevated flagellin-TLR5 mediated NF-κB signaling (Hayashi et al., 2001; Voogdt et al., 2018). Additionally, MB4’s increased mucus degrading capacity is a microbial attribute associated with increased intestinal inflammation (Desai et al., 2016). Unexpectedly, the MB1 isolate--which is non-motile and has WT levels of mucin degrading gene expression and activity—is equally as pro-inflammatory as MB4.

A trait that is common to these two pro-inflammatory isolates is defective MbpA cell-surface regulation, indicating a crucial role for this GlcNAc-responsive adhesin in limiting the inflammatory capacity of Aer01. Consistent with this idea, controlled inhibition of MbpA function in WT Aer01 is sufficient to stimulate intestinal inflammation and displaces Aer01 from the anterior midgut region into the foregut, mimicking traits of the evolved MB1 isolate. We found the PbH1 domain repeat architecture of MbpA in various adhesins encoded by vertebrate intestinal symbionts, including human-associated *Akkermansia* species. Like Aer01, *Akkermansia* increases production of a PbH1-containing adhesin (Amuc_*1620*) in the presence of mucin (Ottman et al., 2017), suggesting a similar role in mucus-mediated aggregation. Consistent with this idea, ectopic Amuc_*1620* expression is sufficient to rescue aggregation defects of MB1 isolate in culture and in the intestine and to reduce its pro-inflammatory activity.

We speculate that MbpA activity could limit Aer01’s inflammatory potential through two potential mechanisms. Aer01 interactions with mucin could shield potentially immunogenic epitopes from host cells, as has been shown for some mucin absorbed bacteria (Mikhalchik et al., 2020). Additionally, MbpA could act on host cells directly to inhibit inflammatory behaviors, as has been shown for a PbH1-containing adhesin from *Streptococcus pneumoniae* that inhibits phagocytosis (Yamaguchi et al., 2019). The impacts of PbHI proteins will be context-dependent and contingent on local mucin-derived cues and bacterial behaviors that they regulate. One PbH1 adhesin identified in our analysis, Fap2 of *Fusobacterium nucleatum*, regulates co-aggregation between *F. nucleatum* and *Porphyromonas gingivalis* (Coppenhagen-Glazer et al., 2015) and binds Gal-GalNAc epitopes which are overexpressed in colon rectal cancer (CRC). Fap2 was found to direct *F. nucleatum* localization to mouse colon tumors (Abed et al., 2016) and promote invasion of cultured HCT116 CRC cells and stimulation of IL-8 and CXCL1, which are associated with CRC progression (Casasanta et al., 2020).

Similar to Fabp2’s role in *F. nucleatum* and *P. gingivalis* co-aggregation, we have shown that MbpA plays a role in bacterial-bacterial interactions in the larval zebrafish intestine. In model bacterial communities in gnotobiotic zebrafish, we found that Aer01 and a highly aggregated zebrafish isolate *Enterobacter* ZOR0014 (Ent14) have strong antagonistic relationships (Sundarraman et al., 2020). In combination with the MB4 variant of Aer01, Ent14 was at an even greater colonization disadvantage than with WT Aer01. The presence of MB4 dramatically alters Ent14’s population structure, causing Ent14 to rapidly disaggregate. This disaggregation behavior may be due to MB4’s altered cell surface chemistry or its increased production of carbohydrate-active enzymes, as many mucin-associated glycans, including GlcNAc, are also components of bacterial extracellular polysaccharides (Bales et al., 2013). MB4 could also indirectly act on Ent14 aggregates through its pro-inflammatory capacity in the host, with disaggregation being triggered by MB4-induced host inflammation.

As microbiome research transitions from descriptive to mechanistic studies, it will be important to identify the most tractable features of host-microbe systems that can be tuned to promote system health. Our study reveals how bacteria sense and respond to a dominant feature of the intestinal environment, mucin glucans, to regulate their cell wall properties and aggregation behavior. Cell surface adhesion expression impacts not only immediate properties of the bacterial cells such as the size of cellular aggregates in which they reside, but also organ-wide features such as bacterial distribution along the length of the gut, bacterial-bacterial competitive interactions, and host inflammatory tone. These system-wide features are altered both by impairment of bacterial mucin sensing and even by disruption of a single feature of their mucin response program, cell surface retention of a mucin-responsive adhesin. These findings point to multiple avenues for manipulating microbiota-member behaviors. As a proof of concept, we demonstrate that expression of a MbpA-like *Akkeromansia* mucin-responsive adhesin in Aer01 is sufficient to reverse its proinflammatory activity. Engineered mucin-responsive cell surface adhesins may be useful in other context to mask proinflammatory bacterial cell surface molecules, aggregate proinflammatory community members, or modulate host immune responses.

## Acknowledgments

We thank Cathy Robinson for her motility assay insight. We thank Rose Sockol and UO Zebrafish Facility staff for expert fish husbandry. We thank Caitlin Kowalski and members of the Guillemin lab for thoughtful discussion throughout this study. Research reported in this publication was supported by the National Institutes of Health under award numbers F32DK124033 and 1P01GM125576. The content is solely the responsibility of the authors and does not necessarily represent the official views of the National Institutes of Health.

## Author Contributions

Conceptualization, T.J.S and K.G.; methodology, T.J.S., D.S., E.M., R.P., and K.G.; formal analysis, T.J.S. and D.S.; writing – original draft T.J.S. and K.G.; writing – review and editing, T.J.S., D.S., R.P., and K.G.; funding acquisition, T.J.S., R.P., and K.G.; resources, R.P. and K.G.

## Declaration of Interests

The authors declare no competing interests.

## Methods

### RESOURCE AVAILABILITY

#### Lead contact

Further information and requestis for resources should be directed to the corresponding author, Karen Guillemin.

#### Materials availability

Bacterial strains and plasmids generated by this study are available through the lead contact.

#### Data and code availability

The raw sequence reads will soon be available through BioProject.

### EXPERIMENTAL MODEL AND SUBJECTS DETAILS

#### Ethics statement, animal care, and gnotobiology

All experiments with zebrafish (*Danio rerio*) were performed according to protocols approved by the University of Oregon Institutional Animal Care and Use Committee and following standard protocols. The zebrafish strains employed in this study were wild type (AB x TU) and Tg(BACmpx:GFP)i114 (referred as MPX:GFP) (Renshaw et al., 2006). Gnotobiotic (germ free) larval zebrafish were derived as previously described (Melancon et al., 2017). Briefly, 0 dfp embryos were washed with sterile embryo medium (SEM) containing antibiotics, then sequentially submerged in bleach and iodine solutions. Sterilized embryos were transferred to tissue culture flasks containing SEM at densities of 1 embryo/mL. The flasks were stored in a temperature controlled room at 28°C.

Prior to all imaging experiments, larve were anesthetized in SEM including 20 μL/mL MS-222. Larvae were euthanized using SEM with 40 μL/mL MS-222.

#### Bacterial strains

For all experiments, bacterial strains were cultured overnight at 30°C in lysogeny broth (LB) medium with shaking. Gentamicin (10μg/mL), kanamycin (50μg/mL), or ampicillin (100 μg/mL) were used when appropriate. When selecting for recombinant Aer01 tagged with the pTn7xTS or pYTn7xTS Tn7 transposon, the triparental conjugation mixture was played on tryptic soy agar (TSA) containing 25 μg/mL gentamicin and 50 μg/mL kanamycin. Tn7-tagged Aer01 were grown in gentamicin. For anhydrotetracycline (aTc) induction, aTc was added at 50 ng/mL in culture or 200 ng/mL to flask water. Oligonucleotides used in this study are provided in Table S2.

##### Generation of mbpA::mNeonGreen reporter strain

To track Aer01 and monitor *mbpA* expression in culture and in the intestine we designed a synthetic construct where dTomato is constitutively expressed (P_tac_::dTom) and mNeonGreen is conditionally expressed by the putative *mbpA* promoter containing a synthetic RBS (*P_mbpA_::mNeonGreen*). The P_tac_::dTom cassette was cloned out of pTW244 using primers JP33 and JP34. The putative *mbpA* promoter (nucleotides −25 to −280 from the *mbpA* ORF start site) with the RBS replaced by a synthetic RBS was cloned from Aer01 genomic DNA using primers JP67/JP68. Primers JP69/JP38 were used to clone mNeonGreen from pSH917::mNeonGreen. Gibson Assembly was used to assemble and insert the full construct into the SmaI-digested Tn7 tagging vector pTn7xTS (Wiles et al., 2018).

##### Generation of aTc-inducible constructs

The Tn7 tagging constructs for inducing *mbpG, mbpD*, and *mucA* expression were modeled after aTc-inducible genetic switches recently engineered by our lab (Wiles et al., 2020). For these constructs, a variant of pTn7xTS that uses native recombination in *Saccharomyces cerevisiae* to combine cloning amplicons was generated (pYTn7xTS) to facilitate assembly of large DNA fragments. Here, the Amp^R^ cassette of pTn7xTS was replaced with an autonomously replicating yeast centromere and URA3 gene for yeast cloning compatibility from the plasmid pMQ30 (Shanks et al., 2006) using primers JP27/JP28 and JP29/JP30. The Tet operator and P_tac_::dTomato-TetR regions were first amplified from pTW324 using the primer pairs JP85/JP86 and JP83/JP84. The *mbpG* ORF was amplified from Aer01 genomic DNA using primer pairs JP165/JP88. The *mbpD* ORF was amplified using primers

The *mucA* ORF was amplified from *Akkermansia muciniphilia* ATCC BAA-835 genomic DNA purchased from ATCC (BAA-835D-5) using primers JP179/JP180. The plasmid pYTn7xTS was digested with SmaI prior to yeast cloning.

##### Triparental mating

Aer01 and MB isolates were tagged with the pTn7xTS or pYTn7xTS constructs as recently described (Wiles et al., 2018). Briefly, triparental conjugation was performed with the *Aeromonas* target strain, an *E. coli* SM10 donor strain carrying the transposase-containing pTNS2 helper plasmid, and an *E. coli* S17 donor strain carrying the pTn7xTS or pYTn7xTS-based tagging construct. To select for Aer01 recombinants, the triparental mating was plated on TSA plates containing gentamycin and kanamycin, as both donor strains are Kan^S^.

### Methods Details

#### Experimental Evolution of Aer01

To select for Aer01 that do not aggregate when exposed to GlcNAc, the aggregation assay was performed as detailed above, but after 6 hr, a sample of the planktonic fraction was expanded in LB for passaging. The following day, a sample of the overnight growth was cryopreserved and passaged again in GlcNAc. A total of 7 passages was performed.

Evolved isolates were purified by plating cryopreserved stocks of whole populations from passages 5 and 6 on TSA plates. After incubating for 18 hr. at 30°C, isolates were randomly picked and grown overnight in TSB with shaking in a 96-well plate. Isolates were cryopreserved in 96-well plates in 25% glycerol and stored at −80°C.

#### Aggregation assay

Overnight cultures of Aer01 were washed 3 times in sterile buffer, then diluted 1:10 in buffer containing 0.4% GlcNAc in a 16-well plate and incubated at 30°C for 6 hr with gentle rotation (115 RPM). To quantify aggregation, a sample of the planktonic fraction containing unaggregated Aer01 cells was taken. The Aer01 aggregates were gently resuspended by pipetting and a sample of the resuspension representing the total bacteria taken. The OD_600_ for the total and planktonic samples were taken and the aggregation index calculated as the ratio of total/planktonic cells.

#### Preparation of genomic DNA for sequencing

Genomic DNA was extracted from overnight cultures using the QIAGEN DNeasy Blood & Tissue Kit (QIAGEN Cat#69506). Library preparation and sequencing were conducted at the Microbial Genome Sequencing Center (MiGS, (https://www.migscenter.com/).

#### Preparation of RNA for sequencing

RNA was harvested from wild type Aer01 grown in buffer supplemented with 0.4% glucose or 0.4% GlcNAc, and MB1 and MB4 grown in buffer supplemented with 0.4% GlcNAc using the RNeasy Qiagen kit (Cat. #74104) after 6 hr of exposure. Sequencing and transcript analysis (transcript level quantification, count normalization, and differential expression analysis) was performed at the Microbial Genome Sequencing Center (MiGS, (https://www.migscenter.com/).

#### Quantification of pNP hydrolysis

Stock solutions of 0.1% pNP-GlcNAc, pNP-Gal, pNP-Fuc, or pNP-GalNAc were diluted 1:1 in buffer solution (SEM) containing stationary Aer01 cells (OD_600_~0.05). pNP hydrolysis (indicated by OD_405_) was monitored for 5 hours in a 96-well plate (100μl total volume). Readings taken every 5 minutes on BMG LABTECH FLUOstar Omega.

#### Swim plate assay

Aer01 swim diameter was assayed as detailed previously (Robinson et al., 2018). Briefly, overnight cultures were grown in LB at 30°C. Each strain normalized washed in buffer (SEM) and 5 μl spotted onto swim agar plates (0.2% agar). For swim analysis in rich medium, LB was used. For swim analysis in minimal medium SEM was used and supplemented with 0.4% GlcNAc or 1mM Proline. LB swim plates were incubated for 6 hr at 30°C and SEM plates were incubated for 24 hr.

#### Confocal microscopy of Aer01 and mucus distribution in fixed larvae

GF larval zebrafish inoculated on 4 dpf with dTomato-tagged WT, MB1 or MB4 Aer01 and colonized larvae were fixed 24 hr later in PBS containing 4% paraformaldehyde (PFA). Larvae were rinsed in PBS, washed in PBS + 0.5% Triton-X, and permealized in deionized water. Larvae were blocked in PBS containing 5% NGS, 2% BSA, 1% DMSO (blocking solution). Aer01 tagged with dTomato were labeled by incubating larvae in blocking solution containing Rabbit anti-dsRed (Takara Bio #6324960) at 1:500 dilution. The primary anti-body was washed away in PBS containing 1% DMSO (wash buffer) and larvae were incubated in blocking solution containing Goat anti-Rabbit^546^ (ThermoFisher #A11010) at 1:1000 dilution. Secondary antibody was washed away in wash buffer. Host mucus was labeled by adding WGA-FITC (Vector Laboratories, #FL-1021) at 1:100 in wash buffer. Samples were mounted in Prolong Diamond antifade (Cat # P36966). Image acquisition was performed using a Leica SPE laser scanning confocal microscope.

#### Analysis of Aer01 population structure by stereomicrscopy

Germ free larval zebrafish were mono-associated on 4 dpf and imaged 24 hr later on 5 dpf. Before imaging, larvae were anesthetized in MS-222 and mounted onto glass slides containing a layer of 4% methylcellulose. A Leica M165 FC Fluorescent Stereo Microscope was used to image of the distribution (ie population structure) of dTomato labeled wild type, MB1, MB2, MB3, and MB4 *Aeromonas* throughout the larval zebrafish gut. The full length of the intestine was manually outlined from the images and the pixel intensity across this region measured using plot profile function in Fiji. The gut length for each fish was normalized to the total pixel distance and the mean intensity at every 1/100 normalized distance calculated. The standard deviation at each position was calculated for each group and plotted with the mean values using the ggplot2 package in R.

#### Live light sheet fluorescence imaging of MB isolates

3D imaging of the larval zebrafish gut was conducted using a previously described home-built set-up (Jemielita et al., 2014). GF larvae were inoculated with dTomato-labeled Aer01, MB1 or MB4 5 dpf and imaged 7dpf using a custom-built light sheet microscope described in detail elsewhere (Jemielita 2014, Mbio). Larvae are anesthetized by placing in MS-222 (Syndel) solution in sterile embryo medium (20μL/mL) for approximately 2 minutes and mounted in 0.6% agarose gel using glass capillaries. The larvae are extruded from the capillaries within a temperature controlled (30°C) sample chamber for imaging. The approximately 1mm larval zebrafish gut is imaged in 3D in 4 sub-regions in ~45s. Lasers at 488 nm and 568nm were used to excite for mNeonGreen and dTomato when indicated. After imaging, larvae are euthanized by placing in MS-222 solution in sterile embryo medium (40μL/mL).

#### Identification of single cells and clusters in 3D light sheet fluorescence images

Single cells and cell clusters were identified as recently described. Briefly, a convolutional neural network was used to classify bacteria and noise blobs (Hay & Parthasarathy, 2018). Potential objects were classified by the network and the output was curated manually. For aggregate segmentation, the zebrafish gut was segmented in 2D, images were threshold and the total number of cells in an aggregate determined by dividing the total fluorescence intensity from the segmented object by the median intensity of individuals in the image. Once the single cell and aggregate net populations were determined, they were used to calculate the planktonic fraction.

#### Aer01 and mucin distribution in culture

Aer01 aggregates were developed as detailed in *aggregation assay* in buffer containing WGA^FITC^-stained mucin and imaged using fluorescence stereomicroscopy. To stain mucin, WGA^FITC^ was added to buffer supplemented with 0.4% Type III Mucin (Sigma, cat #M1778) at 5 μg/mL. The solution was incubated for 15 min at room temperature with gentle rotation then pelleted and washed three times in sterile buffer to remove excess WGA^FITC^.

#### Quantification of intestinal neutrophils

The intestines of tricane-euthanized 5 dfp Tg(MPX:GFP) larval zebrafish were dissected using previously described methods. Fluorescent myeloperoxidase positive (MPX+) were enumerated throughout the entire gut or in the mid and distal intestine.

#### Identification of human-associated microbiota with PbH1 adhesins

The SMART database (http://smart.embl-heidelberg.de/) was queried using the term “PbH1 AND Autotransporter”. The species retrieved from this search were cross-referenced with a list of bacteria species in the IBD TaMMA (https://github.com/Humanitas-Danese-s-omics/ibd-meta-analysis-data/blob/main/manual/bacteria_species_list.tsv).

### QUANTIFICATION AND STATISTICAL ANALYSIS

#### Statistical analysis

Statistical analyses were performed using Prism9 (GraphPad Software). The statistical test used and number of samples are included and described in each figure legend. Group statistics (e.g., mean or median) are indicated in the figure legends when plotted.

## Supplemental Information

**Figure S1. Aer01 associates with WGA-stained mucin in culture.** dTomato-tagged Aer01 were exposed to 0.4% mucin that was stained with WGA (left) or untreated (right) in the aggregation assay. The merged image shows Aer01 (red) associated with WGA-stained mucin (green). WGA-stained mucin does not aggregate unless Aer01 is present (not shown).

**Figure S2. MB4 represses *mbpA* when exposed to GlcNAc.** Quantification of GFP expression in WT, MB1, or MB4 carrying the P*_mbpA_::GFP* construct as a single copy on the chromosome at the *attTn7* site. Reading were taken after 6 hrs in MM supplemented with 0.4% GlcNAc.

**Figure S3. MB1 GlcNAc-mediated aggregation is rescued by *mbpD* expression.** Aggregation assay in GlcNAc with MB1 expressing empty vector or the wild-type *mbpD* allele. Expression of *mbpD* rescues GlcNAc-mediated aggregation of MB1 which contains a LOF mutation in *mbpD*.

**Figure S4. A hyper-motile Aer01 mutant responds to GlcNAc.** A hyper-motile Aer01 mutant, ΔspdE, aggregates when exposed to GlcNAc in the aggregation assay for 6 hr.

**Movie S1. Association between Aer01 and mucus signals in the anterior midgut of fixed intestines.** Scan of z-stacks from region used for the maximum projection image in Fig. 1B.

**Movie S2. Association between Aer01 and mucus signals in the distal midgut of fixed intestines.** Scan of z-stacks from region used for the maximum projection image in Fig. 1C.

**Movie S3. MB1 are planktonic and non-motile in the zebrafish foregut.** Light sheet fluorescence microscopy was used to image a single plane of the zebrafish intestine over time.

**Movie S4. MB4 are planktonic and motile in the zebrafish foregut.** Light sheet fluorescence microscopy was used to image a single plane of the zebrafish intestine over time.

## Notes

### Competing Interest Statement

The authors have declared no competing interest.

## References

Abed, J., Emgård, J. E. M., Zamir, G., Faroja, M., Almogy, G., Grenov, A., Sol, A., Naor, R., Pikarsky, E., Atlan, K. A., Mellul, A., Chaushu, S., Manson, A. L., Earl, A. M., Ou, N., Brennan, C. A., Garrett, W. S., & Bachrach, G. (2016). Fap2 Mediates Fusobacterium nucleatum Colorectal Adenocarcinoma Enrichment by Binding to Tumor-Expressed Gal-GalNAc. Cell Host and Microbe, 20(2), 215–225. https://doi.org/10.1016/j.chom.2016.07.006

Arike, L., Holmén-Larsson, J., & Hansson, G. C. (2017). Intestinal Muc2 mucin O-glycosylation is affected by microbiota and regulated by differential expression of glycosyltranferases. Glycobiology, 27(4), 318–328. https://doi.org/10.1093/glycob/cww134

Azimi, S., Thomas, J., Cleland, S. E., Curtis, J. E., Goldberg, J. B., & Diggle, S. P. (2021). Cell surface hydrophobicity determines aggregate assembly type in Pseudomonas aeruginosa. MBio, 12(4). https://doi.org/https://doi.org/10.1128/mBio.00860-21

Bales, P. M., Renke, E. M., May, S. L., Shen, Y., & Nelson, D. C. (2013). Purification and Characterization of Biofilm-Associated EPS Exopolysaccharides from ESKAPE Organisms and Other Pathogens. PLoS ONE, 8(6). https://doi.org/10.1371/journal.pone.0067950

Banse, A. V, Vanbeuge, S., Smith, T. J., Logan, S. L., & Guillemin, K. (2022). Secreted Aeromonas GlcNAc binding protein GbpA stimulates epithelial cell proliferation in the zebrafish intestine. BioRxiv. https://doi.org/https://doi.org/10.1101/2022.06.27.497793

Bates, J. M., Mittge, E., Kuhlman, J., Baden, K. N., Cheesman, S. E., & Guillemin, K. (2006). Distinct signals from the microbiota promote different aspects of zebrafish gut differentiation. Developmental Biology, 297(2), 374–386. https://doi.org/10.1016/j.ydbio.2006.05.006

Bergstrom, K., Shan, X., Casero, D., Batushansky, A., Lagishetty, V., Jacobs, J. P., Hoover, C., Kondo, Y., Shao, B., Gao, L., Zandberg, W., Noyovitz, B., Michael McDaniel, J., Gibson, D. L., Pakpour, S., Kazemian, N., McGee, S., Houchen, C. W., Rao, C. V., … Xia, L. (2020). Proximal colon-derived O-glycosylated mucus encapsulates and modulates the microbiota. Science, 370(6515), 467–472. https://doi.org/10.1126/science.aay7367

Bergstrom, K., & Xia, L. (2022). The barrier and beyond: Roles of intestinal mucus and mucin-type O-glycosylation in resistance and tolerance defense strategies guiding host-microbe symbiosis. Gut Microbes, 14(1). https://doi.org/10.1080/19490976.2022.2052699

Bhowmick, R., Ghosal, A., Das, B., Koley, H., Saha, D. R., Ganguly, S., Nandy, R. K., Bhadra, R. K., & Chatterjee, N. S. (2008). Intestinal adherence of Vibrio cholerae involves a coordinated interaction between colonization factor GbpA and mucin. Infection and Immunity, 76(11), 4968–4977. https://doi.org/10.1128/IAI.01615-07

Boyd, C. D., Chatterjee, D., Sondermann, H., & O’Toole, G. A. (2012). LapG, required for modulating biofilm formation by pseudomonas fluorescens Pf0-1, is a calcium-dependent protease. Journal of Bacteriology, 194(16), 4406–4414. https://doi.org/10.1128/JB.00642-12

Burns, A. R., & Guillemin, K. (2017). The scales of the zebrafish: host–microbiota interactions from proteins to populations. Current Opinion in Microbiology, 38, 137–141. https://doi.org/10.1016/j.mib.2017.05.011

Casasanta, M. A., Yoo, C. C., Udayasuryan, B., Sanders, B. E., Umanã, A., Zhang, Y., Peng, H., Duncan, A. J., Wang, Y., Li, L., Verbridge, S. S., & Slade, D. J. (2020). Fusobacterium nucleatum host-cell binding and invasion induces IL-8 and CXCL1 secretion that drives colorectal cancer cell migration. Science Signaling, 13(641), 1–13. https://doi.org/10.1126/SCISIGNAL.ABA9157

Co, J. Y., Cárcamo-Oyarce, G., Billings, N., Wheeler, K. M., Grindy, S. C., Holten-Andersen, N., & Ribbeck, K. (2018). Mucins trigger dispersal of Pseudomonas aeruginosa biofilms. Npj Biofilms and Microbiomes, 4(1), 23. https://doi.org/10.1038/s41522-018-0067-0

Collins, A. J., Jarrod Smith, T., Sondermann, H., & O’Toole, G. A. (2020). From Input to Output: The Lap/c-di-GMP Biofilm Regulatory Circuit. Annual Review of Microbiology, 74, 607–631. https://doi.org/10.1146/annurev-micro-011520-094214

Coppenhagen-Glazer, S., Sol, A., Abed, J., Naor, R., Zhang, X., Han, Y. W., & Bachrach, G. (2015). Fap2 of Fusobacterium nucleatum is a galactose-inhibitable adhesin involved in coaggregation, cell adhesion, and preterm birth. Infection and Immunity, 83(3), 1104–1113. https://doi.org/10.1128/IAI.02838-14

Crouch, L. I., Liberato, M. V., Urbanowicz, P. A., Baslé, A., Lamb, C. A., Stewart, C. J., Cooke, K., Doona, M., Needham, S., Brady, R. R., Berrington, J. E., Madunic, K., Wuhrer, M., Chater, P., Pearson, J. P., Glowacki, R., Martens, E. C., Zhang, F., Linhardt, R. J., … Bolam, D. N. (2020). Prominent members of the human gut microbiota express endo-acting O-glycanases to initiate mucin breakdown. Nature Communications, 11(1). https://doi.org/10.1038/s41467-020-17847-5

Cullender, T. C., Chassaing, B., Janzon, A., Kumar, K., Muller, C. E., Werner, J. J., Angenent, L. T., Bell, M. E., Hay, A. G., Peterson, D. A., Walter, J., Vijay-Kumar, M., Gewirtz, A. T., & Ley, R. E. (2013). Innate and adaptive immunity interact to quench microbiome flagellar motility in the gut. Cell Host and Microbe, 14(5), 571–581. https://doi.org/10.1016/j.chom.2013.10.009

Deatherage, D. E., & Barrick, J. E. (2014). Identification of mutations in laboratory evolved microbes from next-generation sequencing data using breseq. Methods Mol Biol., 1151, 165–188. https://doi.org/10.1007/978-1-4939-0554-6_12

Desai, M. S., Seekatz, A. M., Koropatkin, N. M., Kamada, N., Hickey, C. A., Wolter, M., Pudlo, N. A., Kitamoto, S., Terrapon, N., Muller, A., Young, V. B., Henrissat, B., Wilmes, P., Stappenbeck, T. S., Núñez, G., & Martens, E. C. (2016). A Dietary Fiber-Deprived Gut Microbiota Degrades the Colonic Mucus Barrier and Enhances Pathogen Susceptibility. Cell, 167(5), 1339–1353.e21. https://doi.org/10.1016/j.cell.2016.10.043

Earle, K. A., Billings, G., Sigal, M., Lichtman, J. S., Hansson, G. C., Elias, J. E., Amieva, M. R., Huang, K. C., & Sonnenburg, J. L. (2015). Quantitative Imaging of Gut Microbiota Spatial Organization. Cell Host and Microbe, 18(4), 478–488. https://doi.org/10.1016/j.chom.2015.09.002

Flores, E. M., Nguyen, A. T., Odem, M. A., Eisenhoffer, G. T., & Krachler, A. M. (2020). The zebrafish as a model for gastrointestinal tract–microbe interactions. Cellular Microbiology, 22(3). https://doi.org/10.1111/cmi.13152

Frese, S. A., MacKenzie, D. A., Peterson, D. A., Schmaltz, R., Fangman, T., Zhou, Y., Zhang, C., Benson, A. K., Cody, L. A., Mulholland, F., Juge, N., & Walter, J. (2013). Molecular Characterization of Host-Specific Biofilm Formation in a Vertebrate Gut Symbiont. PLoS Genetics, 9(12). https://doi.org/10.1371/journal.pgen.1004057

Grys, T. E., Walters, L. L., & Welch, R. A. (2006). Characterization of the StcE protease activity of Escherichia coli O157:H7. Journal of Bacteriology, 188(13), 4646–4653. https://doi.org/10.1128/JB.01806-05

Hansson, G. C. (2012). Role of mucus layers in gut infection and inflammation. Current Opinion in Microbiology, 15(1), 57–62. https://doi.org/10.1016/j.mib.2011.11.002

Hay, E. A., & Parthasarathy, R. (2018). Performance of convolutional neural networks for identification of bacteria in 3D microscopy datasets. PLoS Computational Biology, 14(12), 1–17. https://doi.org/10.1371/journal.pcbi.1006628

Hayashi, F., Smith, K. D., Ozinsky, A., Hawn, T. R., Yi, E. C., Goodlett, D. R., Eng, J. K., Akira, S., Underhill, D. M., & Aderem, A. (2001). The innate immune response to bacterial flagellin is mediated by Toll-like receptor 5. Nature, 410(6832), 1099–1103. https://doi.org/10.1038/35074106

Hecht, A. L., Casterline, B. W., Choi, V. M., & Bubeck Wardenburg, J. (2017). A Two-Component System Regulates Bacteroides fragilis Toxin to Maintain Intestinal Homeostasis and Prevent Lethal Disease. Cell Host and Microbe, 22(4), 443–448.e5. https://doi.org/10.1016/j.chom.2017.08.007

Hews, C. L., Tran, S. L., Wegmann, U., Brett, B., Walsham, A. D. S., Kavanaugh, D., Ward, N. J., Juge, N., & Schüller, S. (2017). The StcE metalloprotease of enterohaemorrhagic Escherichia coli reduces the inner mucus layer and promotes adherence to human colonic epithelium ex vivo. Cellular Microbiology, 19(6), 1–10. https://doi.org/10.1111/cmi.12717

Jemielita, M., Taormina, M. J., Burns, A. R., Hampton, J. S., Rolig, A. S., Guillemin, K., & Parthasarathy, R. (2014). Spatial and temporal features of the growth of a bacterial species colonizing the zebrafish gut. MBio, 5(6), 1–8. https://doi.org/10.1128/mBio.01751-14

Jevtov, I., Samuelsson, T., Yao, G., Amsterdam, A., & Ribbeck, K. (2014). Zebrafish as a model to study live mucus physiology. Scientific Reports, 4, 1–6. https://doi.org/10.1038/srep06653

Johansson, M. E. V., Holmén Larsson, J. M., & Hansson, G. C. (2011). The two mucus layers of colon are organized by the MUC2 mucin, whereas the outer layer is a legislator of host-microbial interactions. Proceedings of the National Academy of Sciences of the United States of America, 108(SUPPL. 1), 4659–4665. https://doi.org/10.1073/pnas.1006451107

Johansson, M. E. V., Phillipson, M., Petersson, J., Velcich, A., Holm, L., & Hansson, G. C. (2008). The inner of the two Muc2 mucin-dependent mucus layers in colon is devoid of bacteria. Proceedings of the National Academy of Sciences of the United States of America, 105(39), 15064–15069. https://doi.org/10.1073/pnas.0803124105

Jones, C. J., Utada, A., Davis, K. R., Thongsomboon, W., Zamorano Sanchez, D., Banakar, V., Cegelski, L., Wong, G. C. L., & Yildiz, F. H. (2015). C-di-GMP Regulates Motile to Sessile Transition by Modulating MshA Pili Biogenesis and Near-Surface Motility Behavior in Vibrio cholerae. PLoS Pathogens, 11(10), 1–27. https://doi.org/10.1371/journal.ppat.1005068

Juge, N. (2012). Microbial adhesins to gastrointestinal mucus. Trends in Microbiology, 20(1), 30–39. https://doi.org/10.1016/j.tim.2011.10.001

Kavanaugh, N. L., Zhang, A. Q., Nobile, C. J., Johnson, A. D., & Ribbeck, K. (2014). Mucins suppress virulence traits of Candida albicans. MBio, 5(6), 1–8. https://doi.org/10.1128/mBio.01911-14

Kim, J. S., Song, S., Lee, M., Lee, S., Lee, K., & Ha, N. C. (2016). Crystal Structure of a Soluble Fragment of the Membrane Fusion Protein HlyD in a Type i Secretion System of Gram-Negative Bacteria. Structure, 24(3), 477–485. https://doi.org/10.1016/j.str.2015.12.012

Koropatkin, N. M., Cameron, E. A., & Martens, E. C. (2012). How glycan metabolism shapes the human gut microbiota. Nature Reviews Microbiology, 10(5), 323–335. https://doi.org/10.1038/nrmicro2746

Larsson, J. M. H., Karlsson, H., Sjövall, H., & Hansson, G. C. (2009). A complex, but uniform O-glycosylation of the human MUC2 mucin from colonic biopsies analyzed by nanoLC/MSn. Glycobiology. https://doi.org/10.1093/glycob/cwp048

Letunic, I., Khedkar, S., & Bork, P. (2021). SMART: Recent updates, new developments and status in 2020. Nucleic Acids Research, 49(D1), D458–D460. https://doi.org/10.1093/nar/gkaa937

Loose, J. S. M., Forsberg, Z., Fraaije, M. W., Eijsink, V. G. H., & Vaaje-Kolstad, G. (2014). A rapid quantitative activity assay shows that the Vibrio cholerae colonization factor GbpA is an active lytic polysaccharide monooxygenase. FEBS Letters, 588(18), 3435–3440. https://doi.org/10.1016/j.febslet.2014.07.036

Malaker, S. A., Pedram, K., Ferracane, M. J., Bensing, B. A., Krishnan, V., Pett, C., Yu, J., Woods, E. C., Kramer, J. R., Westerlind, U., Dorigo, O., & Bertozzi, C. R. (2019). The mucin-selective protease StcE enables molecular and functional analysis of human cancer-associated mucins. Proceedings of the National Academy of Sciences of the United States of America, 116(15), 7278–7287. https://doi.org/10.1073/pnas.1813020116

Maresca, M., Alatou, R., Pujol, A., Nicoletti, C., Perrier, J., Giardina, T., Simon, G., Méjean, V., & Fons, M. (2021). RadA, a MSCRAMM adhesin of the dominant symbiote Ruminococcus gnavus e1, binds human immunoglobulins and intestinal mucins. Biomolecules, 11(11). https://doi.org/10.3390/biom11111613

Massaquoi, M. S., Kong, G., Chilin, D., Hamilton, M. K., & Melancon, E. (2022). GLOBAL HOST RESPONSES TO THE MICROBIOTA AT SINGLE CELL Keywords. https://doi.org/https://doi.org/10.1101/2022.03.28.486083

Massimino, L., Lamparelli, L. A., Houshyar, Y., D’Alessio, S., Peyrin-Biroulet, L., Vetrano, S., Danese, S., & Ungaro, F. (2021). The Inflammatory Bowel Disease Transcriptome and Metatranscriptome Meta-Analysis (IBD TaMMA) framework. Nature Computational Science, 1(8), 511–515. https://doi.org/10.1038/s43588-021-00114-y

Melancon, E., Gomez De La Torre Canny, S., Sichel, S., Kelly, M., Wiles, T. J., Rawls, J. F., Eisen, J., & Guillemin, K. (2017). Best practices for germ-free derivation and gnotobiotic zebrafish husbandry. Methods Cell Biol., 138, 61–100. https://doi.org/10.1016/bs.mcb.2016.11.005

Michaels, L. A., Singh, P. K., Jennings, L. K., Secor, P. R., & Ratjen, A. (2018). Entropically driven aggregation of bacteria by host polymers promotes antibiotic tolerance in Pseudomonas aeruginosa. Proceedings of the National Academy of Sciences, 115(42), 10780–10785. https://doi.org/10.1073/pnas.1806005115

Mikhalchik, E., Balabushevich, N., Vakhrusheva, T., Sokolov, A., Baykova, J., Rakitina, D., Scherbakov, P., Gusev, S., Gusev, A., Kharaeva, Z., Bukato, O., & Pobeguts, O. (2020). Mucin adsorbed by E. coli can affect neutrophil activation in vitro. FEBS Open Bio, 10(2), 180–196. https://doi.org/10.1002/2211-5463.12770

Mondal, M., Nag, D., Koley, H., Saha, D. R., & Chatterjee, N. S. (2014). The Vibrio cholerae extracellular chitinase ChiA2 is important for survival and pathogenesis in the host intestine. PLoS ONE, 9(9). https://doi.org/10.1371/journal.pone.0103119

Ndeh, D., & Gilbert, H. J. (2018). Biochemistry of complex glycan depolymerisation by the human gut microbiota. FEMS Microbiology Reviews, 42(2), 146–164. https://doi.org/10.1093/femsre/fuy002

Newell, P. D., Boyd, C. D., Sondermann, H., & O’Toole, G. A. (2011). A c-di-GMP effector system controls cell adhesion by inside-out signaling and surface protein cleavage. PLoS Biology, 9(2). https://doi.org/10.1371/journal.pbio.1000587

Ng, A. N. Y., De Jong-Curtain, T. A., Mawdsley, D. J., White, S. J., Shin, J., Appel, B., Dong, P. D. S., Stainier, D. Y. R., & Heath, J. K. (2005). Formation of the digestive system in zebrafish: III. Intestinal epithelium morphogenesis. Developmental Biology, 286(1), 114–135. https://doi.org/10.1016/j.ydbio.2005.07.013

Ottman, N., Davis, M., Suarez-Diez, M., Boeren, S., Schaap, P. J., Martins dos Santos, V. A. P., Smidt, H., Belzer, C., & de Vos, W. M. (2017). Genome-Scale Model and Omics Analysis of Metabolic Capacities of Akkermansia muciniphila Reveal a Preferential Mucin-Degrading Lifestyle. Applies and Environmental Microbiology, 83(18), e01014–17.

Ottman, N., HuuskonenL., L., Reunanen, J., Boeren, S., Klievink, J., Smidt, H., Belzer, C., & De Vos, W. M. (2016). Characterization of outer membrane proteome of akkermansia muciniphila reveals sets of novel proteins exposed to the human intestine. Frontiers in Microbiology, 7(JUL), 1–13. https://doi.org/10.3389/fmicb.2016.01157

Pacheco, A. R., Munera, D., Waldor, M. K., Sperandio, V., & Ritchie, J. M. (2012). Fucose sensing regulates bacterial intestinal colonization. Nature, 492(7427), 113–117. https://doi.org/10.1038/nature11623

Pickard, J. M., Maurice, C. F., Kinnebrew, M. A., Abt, M. C., Schenten, D., Golovkina, T. V., Bogatyrev, S. R., Ismagilov, R. F., Pamer, E. G., Turnbaugh, P. J., & Chervonsky, A. V. (2014). Rapid fucosylation of intestinal epithelium sustains host-commensal symbiosis in sickness. Nature, 514(7524), 638–641. https://doi.org/10.1038/nature13823

Renshaw, S. A., Loynes, C. A., Trushell, D. M. I., Elworthy, S., Ingham, P. W., & Whyte, M. K. B. (2006). A transgenic zebrafish model of neutrophilic inflammation. Blood, 108(13), 3976–3978. https://doi.org/10.1182/blood-2006-05-024075

Robinson, C. D., Klein, H. S., Murphy, K. D., Parthasarathy, R., Guillemin, K., & Bohannan, B. J. M. (2018). Experimental bacterial adaptation to the zebrafish gut reveals a primary role for immigration. PLoS Biology, 16(12), 1–26. https://doi.org/10.1371/journal.pbio.2006893

Robinson, C. D., Sweeney, E. G., Ngo, J., Ma, E., Perkins, A., Smith, T. J., Fernandez, N. L., Waters, C. M., Remington, S. J., Bohannan, B. J. M., & Guillemin, K. (2021). Host-emitted amino acid cues regulate bacterial chemokinesis to enhance colonization. Cell Host & Microbe, 29(8), 1221–1234.e8. https://doi.org/10.1016/j.chom.2021.06.003

Schlomann, B. H., & Parthasarathy, R. (2021). Gut bacterial aggregates as living gels. ELife, 10, 1–22. https://doi.org/10.7554/eLife.71105

Schlomann, B. H., Wiles, T. J., Wall, E. S., Guillemin, K., & Parthasarathy, R. (2018). Bacterial Cohesion Predicts Spatial Distribution in the Larval Zebrafish Intestine. Biophysical Journal, 115(11), 2271–2277. https://doi.org/10.1016/j.bpj.2018.10.017

Schlomann, B. H., Wiles, T. J., Wall, E. S., Guillemin, K., & Parthasarathy, R. (2019). Sublethal antibiotics collapse gut bacterial populations by enhancing aggregation and expulsion. Proceedings of the National Academy of Sciences of the United States of America, 116(43), 21392–21400. https://doi.org/10.1073/pnas.1907567116

Shanks, R. M. Q., Caiazza, N. C., Hinsa, S. M., Toutain, C. M., & O’Toole, G. a. (2006). Saccharomyces cerevisiae-based molecular tool kit for manipulation of genes from gram-negative bacteria. Applied and Environmental Microbiology, 72(7), 5027–5036. https://doi.org/10.1128/AEM.00682-06

Sonnenburg, J. L., Xu, J., Leip, D. D., Chen, C. H., Westover, B. P., Weatherford, J., Buhler, J. D., & Gordon, J. I. (2005). Glycan foraging in vivo by an intestine-adapted bacterial symbiont. Science, 307(5717), 1955–1959. https://doi.org/10.1126/science.1109051

Sudarsan, N., Lee, E. R., Weinberg, Z., Moy, R. H., Kim, J. N., Link, K. H., & Breaker, R. R. (2008). Riboswitches in eubacteria sense the second messenger cyclic Di-GMP. Science, 321(5887), 411–413. https://doi.org/10.1126/science.1159519

Sundarraman, D., Hay, E. A., Martins, D. M., Shields, D. S., Pettinari, N. L., & Parthasarathy, R. (2020). Higher-order interactions dampen pairwise competition in the zebrafish gut microbiome. MBio, 11(5), 1–15. https://doi.org/10.1128/mBio.01667-20

Troll, J. V., Hamilton, M. K., Abel, M. L., Ganz, J., Bates, J. M., Stephens, W. Z., Melancon, E., van der Vaart, M., Meijer, A. H., Distel, M., Eisen, J. S., & Guillemin, K. (2018). Microbiota promote secretory cell determination in the intestinal epithelium by modulating host Notch signaling. Development, 145(4), dev155317. https://doi.org/10.1242/dev.155317

Vaishnava, S., Yamamoto, M., Severson, K. M., Ruhn, K. a, Yu, X., Koren, O., Ley, R., Wakeland, E. K., & Hooper, L. V. (2011). The Antibacterial Lectin RegIII-gamma Promotes the Spatial Segregation of Microbiota and Host in the Intestine. Science, 334(October), 255–258.

Voogdt, C. G. P., Wagenaar, J. A., & van Putten, J. P. M. (2018). Duplicated TLR5 of zebrafish functions as a heterodimeric receptor. Proceedings of the National Academy of Sciences, 115(14), E3221–E3229. https://doi.org/10.1073/pnas.1719245115

Wallace, K. N., Akhter, S., Smith, E. M., Lorent, K., & Pack, M. (2005). Intestinal growth and differentiation in zebrafish. Mechanisms of Development, 122(2), 157–173. https://doi.org/10.1016/j.mod.2004.10.009

Wang, B. X., Wheeler, K. M., Cady, K. C., Lehoux, S., Cummings, R. D., Laub, M. T., & Ribbeck, K. (2020). Mucin Glycans Signal through the Sensor Kinase RetS to Inhibit Virulence-Associated Traits in Pseudomonas aeruginosa. Current Biology, 1–13. https://doi.org/10.1016/j.cub.2020.09.088

Wardman, J. F., Bains, R. K., Rahfeld, P., & Withers, S. G. (2022). Carbohydrate-active enzymes (CAZymes) in the gut microbiome. Nature Reviews Microbiology, 0123456789. https://doi.org/10.1038/s41579-022-00712-1

Welch, J. L. M., Hasegawa, Y., McNulty, N. P., Gordon, J. I., & Borisy, G. G. (2017). Spatial organization of a model 15-member human gut microbiota established in gnotobiotic mice. Proceedings of the National Academy of Sciences of the United States of America, 114(43), E9105–E9114. https://doi.org/10.1073/pnas.1711596114

Wheeler, K. M., Cárcamo-Oyarce, G., Turner, B. S., Dellos-Nolan, S., Co, J. Y., Lehoux, S., Cummings, R. D., Wozniak, D. J., & Ribbeck, K. (2019). Mucin glycans attenuate the virulence of Pseudomonas aeruginosa in infection. Nature Microbiology, 4(12), 2146–2154. https://doi.org/10.1038/s41564-019-0581-8

Wiles, T. J., Jemielita, M., Baker, R. P., Schlomann, B. H., Logan, S. L., Ganz, J., Melancon, E., Eisen, J. S., Guillemin, K., & Parthasarathy, R. (2016). Host Gut Motility Promotes Competitive Exclusion within a Model Intestinal Microbiota. PLoS Biology, 14(7), 1–24. https://doi.org/10.1371/journal.pbio.1002517

Wiles, T. J., Schlomann, B. H., Wall, E. S., Betancourt, R., Parthasarathy, R., & Guillemin, K. (2020). Swimming motility of a gut bacterial symbiont promotes resistance to intestinal expulsion and enhances inflammation. In PLoS Biology (Vol. 18, Issue 3). https://doi.org/10.1371/journal.pbio.3000661

Wiles, T. J., Wall, E. S., Schlomann, B. H., Guillemin, K., Parthasarathy, R., Hay, E. A., & Wiles, T. J. (2018). Modernized Tools for Streamlined Genetic Manipulation and Comparative Study of Wild and Diverse Proteobacterial Lineages. MBio, 9(5), 1–19. https://doi.org/10.1128/mbio.01877-18

Willms, R. J., Ocampo Jones, L., Hocking, J. C., & Foley, E. (2021). A Cell Atlas of Microbe-Responsive Processes in the Zebrafish Intestine. SSRN Electronic Journal, 38(5), 110311. https://doi.org/10.2139/ssrn.3778935

Yamaguchi, M., Hirose, Y., Takemura, M., Ono, M., Sumitomo, T., Nakata, M., Terao, Y., & Kawabata, S. (2019). Streptococcus pneumoniae Evades Host Cell Phagocytosis and Limits Host Mortality Through Its Cell Wall Anchoring Protein PfbA. Frontiers in Cellular and Infection Microbiology, 9(August), 1–14. https://doi.org/10.3389/fcimb.2019.00301

