## Supplemental Figures for "A mucin-regulated adhesin determines the intestinal biogeography and inflammatory character of a bacterial symbiont"

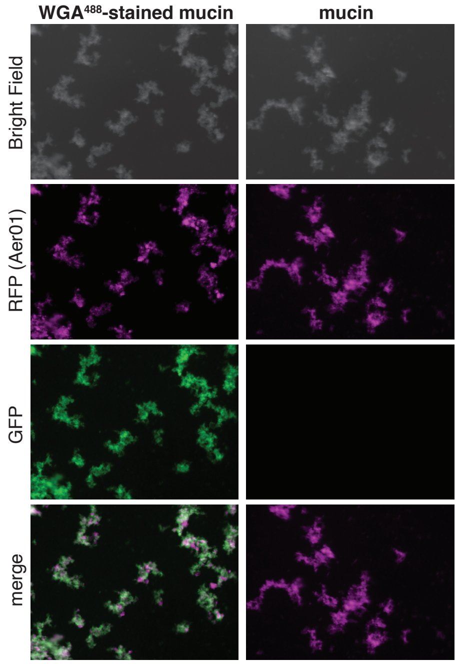


**Figure S1. Aer01 associates with WGA-stained mucin in culture.** dTomato-tagged Aer01 were exposed to 0.4% mucin that was stained with WGA (left) or untreated (right) in the aggregation assay. The merged image shows Aer01 (red) associated with WGA-stained mucin (green). WGA-stained mucin does not aggregate unless Aer01 is present (not shown).

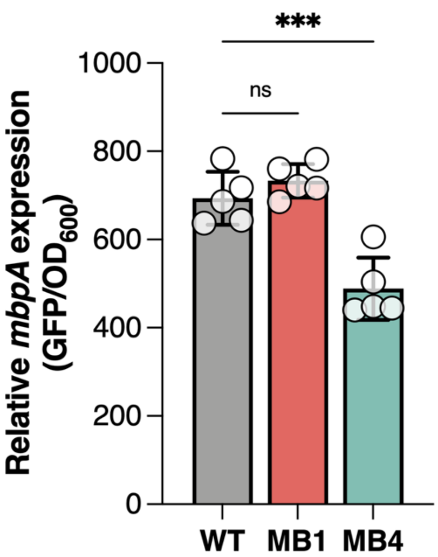


**Figure S2. MB4 represses *mbpA* when exposed to GlcNAc.** Quantification of GFP expression in WT, MB1, or MB4 carrying the $P_{mbpA}$*::GFP* construct as a single copy on the chromosome at the *attTn7* site. Reading were taken after 6 hrs in MM supplemented with 0.4% GlcNAc.

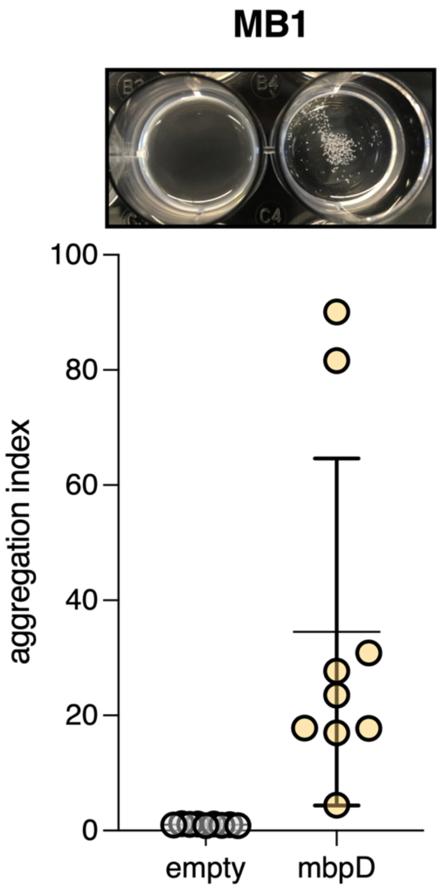


**Figure S3. MB1 GlcNAc-mediated aggregation is rescued by *mbpD* expression.** Aggregation assay in GlcNAc with MB1 expressing empty vector or the wild-type *mbpD* allele. Expression of *mbpD* rescues GlcNAc-mediated aggregation of MB1 which contains a LOF mutation in *mbpD*.

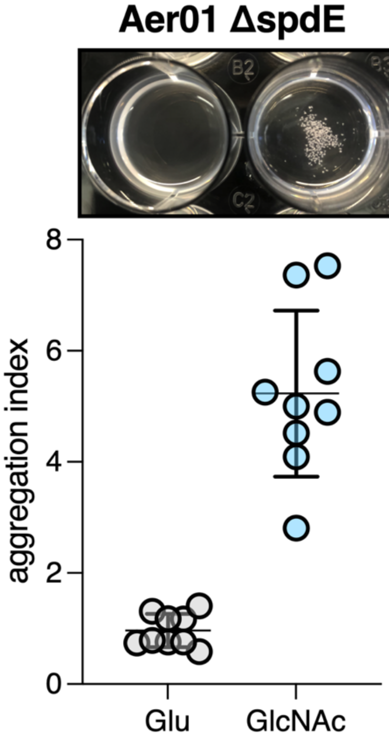


**Figure S4. A hyper-motile Aer01 mutant responds to GlcNAc.** A hyper-motile Aer01 mutant, ΔspdE, aggregates when exposed to GlcNAc in the aggregation assay for 6 hr.
